# Genomes from 117 vertebrate species reveal rapidly evolving segmental duplication landscapes

**DOI:** 10.1101/2024.11.22.624925

**Authors:** Alber Aqil, Saiful Islam, Faraz Hach, Ibrahim Numanagić, Naoki Masuda, Omer Gokcumen

## Abstract

Segmental duplications are major drivers of evolutionary innovation, yet their dynamics across vertebrates remain poorly understood. Here, we identify segmental duplications from long-read sequenced genomes of 117 vertebrates and the starfish, generating the largest multi-species dataset of its kind. We find that vertebrate genomes show a higher propensity for tandem duplications than for interspersed duplications. However, when focusing only on subtelomeric regions, avian and mammalian genomes show the opposite propensity toward interspersed duplications. We also observe that, across vertebrates, tandem duplications tend to be larger than interspersed duplications. Next, we construct a segmental duplication network for each species, and use network-derived properties to quantify the duplication landscape for that species. Functional enrichment analysis of hyper-duplicated genes reveals a strong enrichment in platypus for pheromone response, driven by the expansion of the vomeronasal pheromone receptor *V1R* gene family. Overall, our results uncover the general properties of vertebrate segmental duplication, demonstrate the rapid evolution of segmental duplication landscapes, and highlight the utility of network-based approaches for studying genome evolution.

**Significance:** Gene and regulatory region duplications are a fundamental source of evolutionary raw material. Here we generate segmental duplication calls from 117 vertebrate species. We find that vertebrate genomes have a bias towards tandem duplications relative to interspersed duplications. However, in the subtelomeric regions, birds and mammals exhibit an opposite bias toward interspersed duplications. Our analysis of segmental duplication networks demonstrate that duplication landscapes evolve rapidly, following species-specific rather than phylogenetic patterns. These findings indicate that the genomic architecture underlying segmental duplications is highly dynamic, uniquely shaping each lineage’s potential to adapt. Our study provides the most comprehensive view of vertebrate segmental duplications to date and establishes a network-based framework for studying genomic structural evolution.

## Introduction

Duplication of genes and regulatory regions has been proposed to play a major role in vertebrate evolution (Ohno 1970). In particular, segmental duplications can create redundant paralogs of functional regions (Carroll 2005), allowing mutations that would have been deleterious in the ancestral copy to be tolerated. Such duplications can therefore circumvent valleys in the fitness landscape (Banse et al. 2024) and increase the probability of reaching higher adaptive peaks (Li and Zhang 2025) by enabling the co-option of duplicates for new functions (Innan and Kondrashov 2010). The mutation rate is elevated in segmental duplicates relative to single-copy regions (Vollger et al. 2023), further raising the likelihood that duplication will acquire new functions. Alternatively, duplication of genes or regulatory elements can drive dramatic changes in expression levels (Aqil et al. 2025), which may itself confer a fitness advantage.

Additionally, genomic regions rich in segmental duplication are prone to accumulating further duplications by increased rates of recombination errors (Kim et al. 2008; Lin and Gokcumen 2019), thereby affecting the potential for adaptation. Finally, as functional analyses extend to non-human species, identifying segmental duplications becomes important, since they can also generate spurious functional associations between genomic loci (Aqil and Gokcumen 2025). For these reasons, identifying segmental duplications and studying the evolution of segmental duplication landscapes across diverse species is imperative.

One way to study the evolution of segmental duplication landscapes is through segmental duplication networks. Previous work has shown that models involving gene duplication followed by divergence between duplicates yield realistic properties in biological networks (Solé et al. 2002; Pastor-Satorras et al. 2003; Vázquez et al. 2003; Lin and Zhang 2010; Lü and Wang 2020; Abdullaev et al. 2021). Although duplication-and-divergence network models were first proposed more than two decades ago (Solé et al. 2002; Pastor-Satorras et al. 2003; Vázquez et al. 2003), they have rarely been tested on large empirical datasets of segmental duplications (Bergman et al. 2006; Vollger et al. 2019; Abdullaev et al. 2021; Abdullaev et al. 2023). Such empirical networks can reveal aspects of the segmental duplication landscape, such as the presence of duplication clusters, that non-network approaches cannot capture.

Here, we call genome-wide (including both genic and non-genic regions) segmental duplications from 117 vertebrate species and the starfish using long-read-sequenced genomes (**Figure 1A– B**), generating the largest multi-species catalog of segmental duplications to date. Using this dataset, we explore the properties of vertebrate segmental duplications, and construct segmental duplication networks to study their evolutionary dynamics (**Figure 1C**).

**Figure 1.**
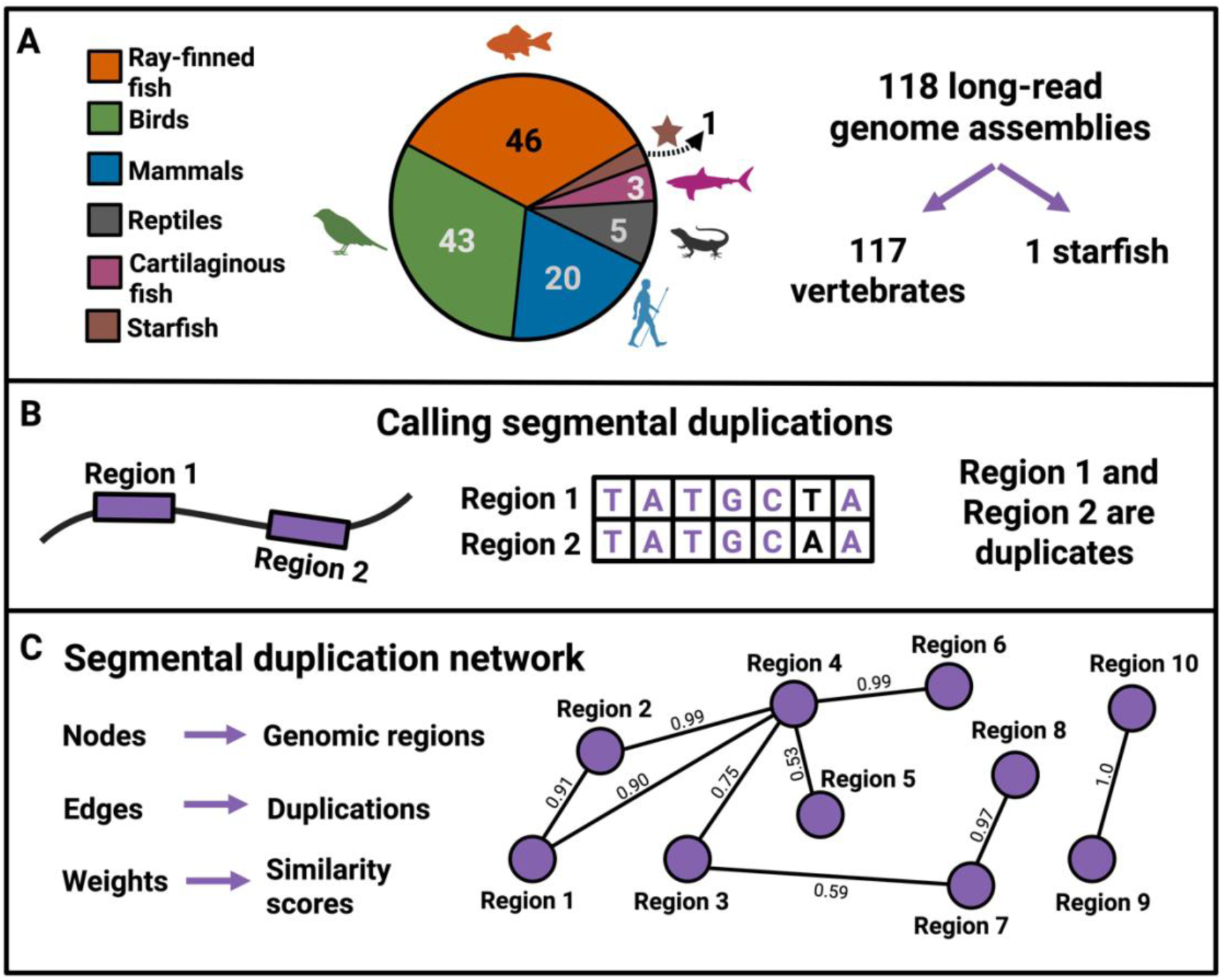
Methodological framework. **A.** The number of species in each taxonomic class in our data. **B.** A schematic showing segmental duplication calling from genome sequences. The image on the left represents a chromosome with two genomic regions marked as Region 1 and Region 2. The middle image shows that the sequences from the two regions align with a high similarity score. Regions 1 and 2 are, therefore, duplicates. **C.** A schematic segmental duplication network. For each species, we used 13 network properties and the average similarity score across duplications (a non-network property) to quantify the segmental duplication landscape.

Overall, our analyses of segmental duplication networks offer new insights into the forces shaping genomic structural variation across vertebrates. By integrating large-scale comparative genomics with network approaches, we address gaps in understanding how the segmental duplication landscape evolves and drives species-specific adaptive potential.

## Results

### Segmental duplication, transposable elements, or ohnologs?

Our approach to detecting segmental duplications relies on identifying regions of sequence similarity with an alignment span of at least 900 bases. Without prior filtering, this approach would capture three categories of duplicated sequence: transposable elements, ohnologs that persist from ancient whole-genome duplications, and segmental duplications.

Transposable elements and short repeats are pervasive in vertebrate genomes (Elliott and Gregory 2015). To remove transposable elements and short repeats, we masked each genome with RepeatMasker using species-specific libraries wherever available. In particular, we used species-specific libraries for all mammals, all birds, all reptiles, 23 out of 46 ray-finned fish species, and 2 out of 3 cartilaginous fish species. For the remaining species, we used class-level repeat libraries. The percentages of each genome masked across taxonomic classes is shown in **Figure 2A**.

**Figure 2.**
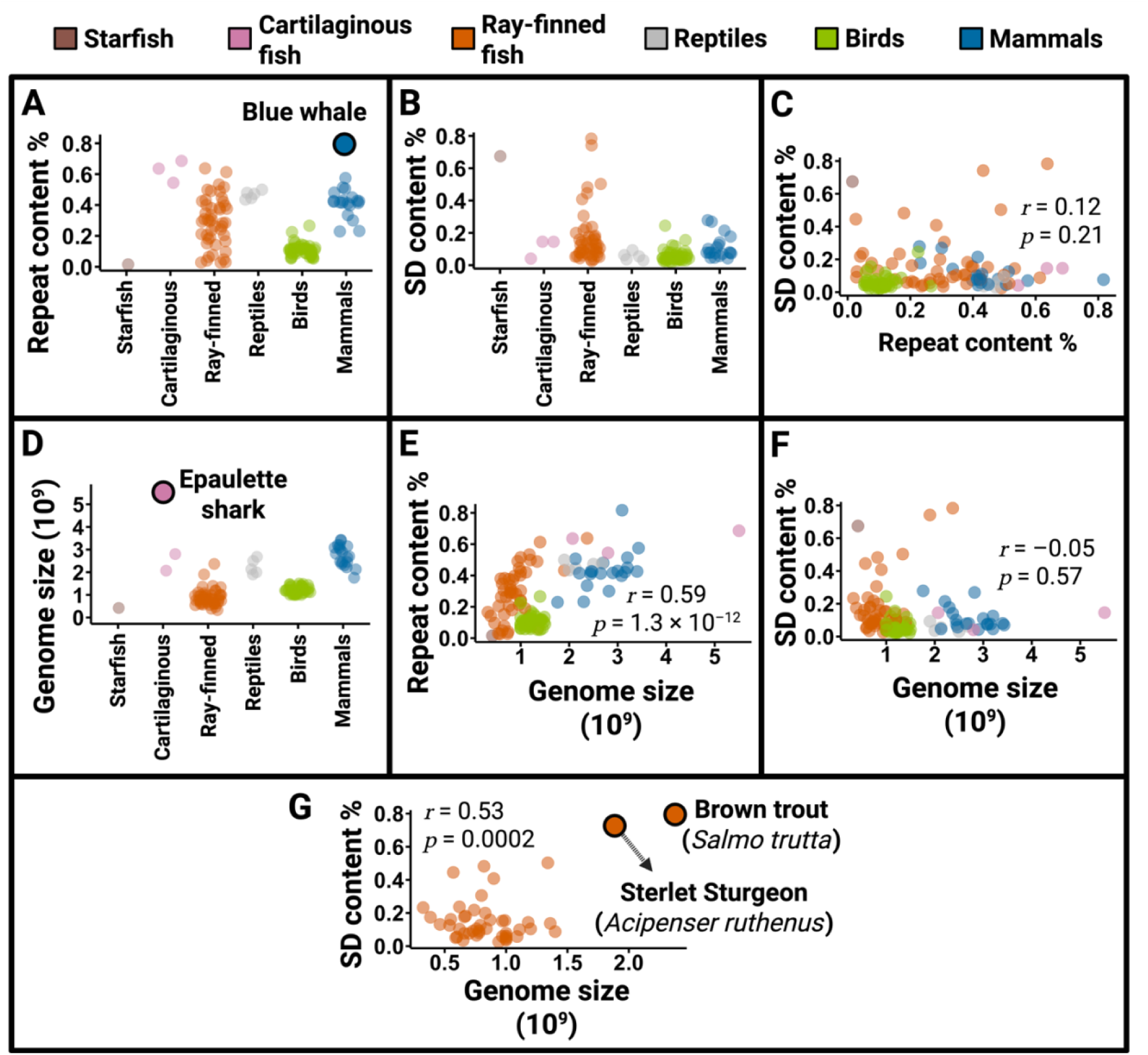
Segmental duplications, repeat content, and ohnologs. **A.** Distribution of the percentage of the genome covered by repeats (transposable elements and simple repeats) across species. The blue whale (*Balaenoptera musculus*) is an outlier among mammals for its high repeat content. **B.** Percentage of the genome covered by segmental duplications across species. **C.** Repeat content and segmental duplication content are uncorrelated across species. **D.** Distribution of genome size across species. The epaulette shark (*Hemiscyllium ocellatum*) is an outlier among our sample of vertebrates for its large genome size (>5 billion base pairs). **E.** Genome size is correlated with repeat content (presumably driven by transposable elements) across species. **F.** Genome size and segmental duplication content are uncorrelated. **G.** The correlation between genome size and the fraction of the genome covered by duplications among ray-finned fish is driven by two outliers: the sterlet sturgeon and the brown trout (a salmonid). After removing these two species, the correlation coefficient drops to –0.05. The apparently high segmental duplication content of brown trout and sterlet sturgeon actually represents a high ohnolog content.

In contrast, we do not expect ohnologs to affect large fractions of genomes. In particular, only two species in our dataset (brown trout and sterlet sturgeon; discussed below) have experienced relatively recent whole-genome duplications. Further, duplicates from ancient whole genome duplications (such as two rounds of genome duplications in ancestral vertebrates) are not detectable due to gene loss and sequence divergence after the whole genome duplication (Langham et al. 2004; Brunet et al. 2006; Cheng et al. 2018), i.e. genomic fractionation, as presciently predicted by Haldane (Haldane 1933). Therefore, with the exception of the brown trout and sterlet sturgeon, ohnologs are unlikely to skew our genome-wide analysis of segmental duplications, and no filter was applied to remove them.

After masking genomes for repeats and transposable elements and reasoning that ohnologs are unlikely to bias our results, we called segmental duplications using BISER (Išerić et al. 2022). The percentage of each genome covered by segmental duplications is shown in **Figure 2B**. These percentages are generally higher than those reported elsewhere because BISER relies on a segmental duplication error model that assumes up to 25% sequence dissimilarity rather than the standard 10%. This allows us to discover older segmental duplications. We note that the repeat content and segmental duplication content across genomes are uncorrelated, suggesting that differential masking quality does not introduce a systematic error in segmental duplication calls (**Figure 2C**).

Next, we examined the potential determinants of genome sizes across species. Consistent with previous work (Hughes and Hughes 1995; Gregory et al. 2009; Kapusta et al. 2017), we find that bird genomes are significantly smaller than those of other amniotes (Cohen’s 𝑑 = 4.5; 𝑝 < 2.2 × 10^−16^; **Figure 2D**). We find that while repeat content, a rough proxy for transposable elements content (Wang et al. 2025), strongly predicts the size of genomes across species (**Figure 2E**), consistent with previous findings (Elliott and Gregory 2015; Osmanski et al. 2023), segmental duplications do not (**Figure 2F**).

However, the sterlet sturgeon and the brown trout, stand out among ray-finned with unusually large genomes and high apparent coverage by segmental duplications (**Figure 2G**). These signals likely reflect the persistence of ohnologs from whole-genome duplications rather than an excess of true segmental duplications, as discussed below.

The sterlet sturgeon (*Acipenser ruthenus*) is a member of the non-teleost “living fossil” sturgeon family. Its lineage has undergone a distinct third round of whole genome duplication (Acipenseriformes-specific third-round or As3R WGD) after divergence from the teleost fish and sometime before 200 million years ago (Redmond et al. 2023). Mirroring its morphological stasis, the sterlet sturgeon’s molecular evolution (Krieger and Fuerst 2002; Du et al. 2020; Redmond et al. 2023), in terms of both substitutions and rearrangements, is so remarkably slow that the duplication event was initially misdated to 21.3 million years ago (Cheng et al. 2019). Indeed, the sterlet sturgeon retains an exceptionally high proportion (70%) of duplicated genes from the As3R event (Bi et al. 2021). This combination of ancient whole genome duplication, high gene retention, and slow evolutionary change likely drives the apparent association between segmental duplication and the sterlet sturgeon’s large genome size.

The brown trout (*Salmo trutta*), a teleost in the salmonid family, is another notable exception. This species, introduced worldwide as a game fish, is recognized as one of the most invasive fish species globally (Lowe et al. 2000). In addition to the teleost-specific whole-genome duplication (Ts3R WGD), the salmonid family underwent a fourth, salmonid-specific genome duplication (Ss4R) between 50 and 80 million years ago (Alexandrou et al. 2013; Macqueen and Johnston 2014; Lien et al. 2016). This extra round of duplication is associated with larger genome sizes in salmonids (Zhu et al. 2022), and as the only salmonid in our dataset, the brown trout has the largest genome among the ray-finned fish we analyzed. Moreover, ∼10-15% of salmonid genomes still exhibit tetrasomic inheritance (Allendorf et al. 2015), promoting gene conversion in these regions (Campbell et al. 2019). These gene conversion events help maintain sequence similarity among duplicated regions, preserving the remnants of the Ss4R event, detectable as spurious segmental duplicates. Thus, the combination of a relatively recent whole-genome duplication and ongoing gene conversion in tetrasomically inherited regions likely explains the brown trout’s large genome and its high fraction of apparent segmental duplications.

### Duplications close together and duplications far away

We find that vertebrate segmental duplications are not distributed randomly across genomes. In particular, a genomic region on a given chromosome is more likely to be duplicated on the same chromosome than on a different chromosome (**Figure 3A**). This is not to say that every genome contains more intrachromosomal duplications in absolute number. Interchromosomal events are more numerous simply because there are many possible non-homologous chromosome pairs. However, when corrected for this larger search space, all vertebrates have a higher density for intrachromosomal relative to interchromosomal duplications. Moreover, even within the same chromosome, a region is more likely to be duplicated in tandem (within 1MB) than farther away (**Figure 3B, Figure S1**). One consequence of this higher propensity for tandem duplications is the formation of tandem arrays of related genes (Pan and Zhang 2008).

**Figure 3.**
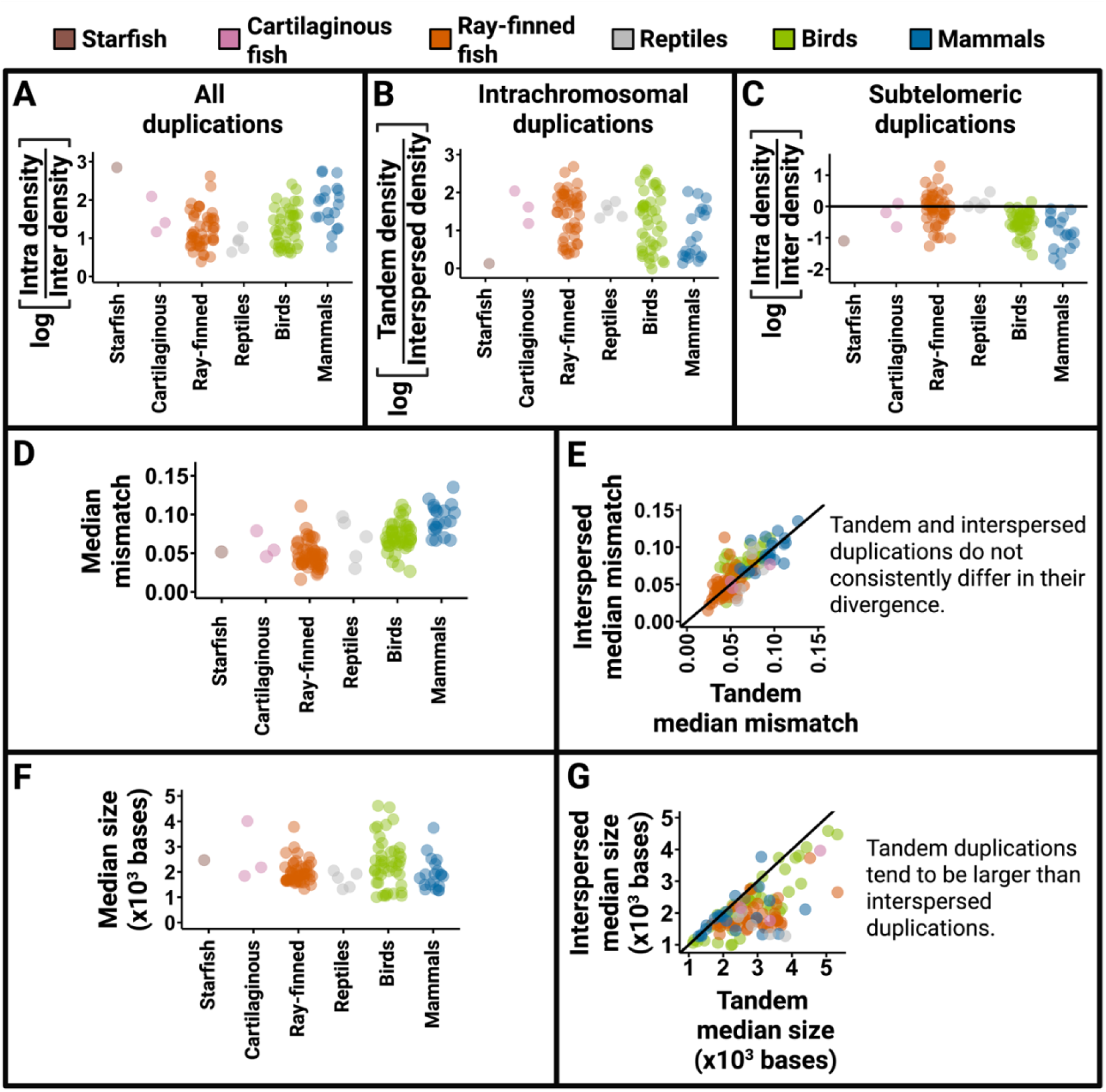
Tandem versus interspersed segmental duplications. **A**. All vertebrate species show a propensity toward intrachromosomal segmental duplications. The y-axis is the logarithm (base 10) of intrachromosomal to interchromosomal duplication densities. A y-axis value > 0 implies intrachromosomal bias. **B.** Intrachromosomal segmental duplications are biased toward tandem (<1 MB apart) as opposed to interspersed (>1 MB apart) events. The y-axis is the logarithm (base 10) of the ratio of tandem to interspersed duplication density. A y-axis value > 0 implies a bias toward tandem duplications. **C.** Subtelomeric duplications in birds, mammals, and approximately half of ray-finned fish species show a propensity toward interchromosomal events. The y-axis is the logarithm (base 10) of the ratio of intrachromosomal to interchromosomal subtelomeric duplication density. A y-axis value > 0 implies an intrachromosomal bias, and a value < 0 implies an interchromosomal bias. **D.** Distribution of median divergence (mismatch score) between partner duplicates across species in different taxonomic classes. **E.** Comparison of median mismatch scores between tandem and interspersed duplications. The diagonal line represents equal mismatch scores of tandem and interspersed duplications. There is no consistent difference between tandem and interspersed mismatch scores. **F.** Distribution of median segmental duplication size across species in different taxonomic classes. **G.** Comparison of median duplication sizes between tandem and interspersed duplications. The diagonal line represents equal median sizes of tandem and interspersed duplications. Species to the right of this line tend to have larger tandem duplications relative to interspersed duplications. Most species (110/118) have larger tandem duplications relative to interspersed duplications.

Interestingly, this density bias toward proximal duplications is reversed near chromosome ends (subtelomeric duplications). Subtelomeric regions in mammals, birds, and about half of ray-finned fish species show a density bias towards interchromosomal duplications (**Figure 3C**). This result mirrors previous findings in humans showing that subtelomeric regions are a pastiche of interchromosomal segmental duplications formed by repeated translocations between chromosome ends, facilitated by non-homologous end joining (Linardopoulou et al. 2005).

These spatial patterns suggest that two broad mechanisms shape duplication landscapes. Local errors in recombination and DNA replication create a propensity toward tandem duplications, whereas non-homologous end joining creates a subtelomeric propensity toward interchromosomal duplications.

Finally, we examined how the size (alignment span) and sequence divergence (mismatch score) of duplications differ between tandem (<1 Mb apart) and interspersed (>1 Mb or interchromosomal) duplicates. We find that the median divergence between duplicates is low (**Figure 3D; Figure S2**), and there is no consistent difference in divergence between tandem and interspersed duplications across species (**Figure 3E**).

More interestingly, we observe that while most duplications are small, on the order of 10^3^ bp (**Figure 3F; Figure S3**), tandem duplications tend to be larger than interspersed ones (**Figure 3G**). This size difference is consistent with observations in humans (Zhang et al. 2005), mice (Francoeur et al. 2025), and even drosophila (Fiston-Lavier et al. 2007), suggesting that it may be a general feature of animal genomes. This result may also explain the finding in mammals that interchromosomal daughter gene copies evolve faster relative to their parent paralog than do intrachromosomal daughter copies (Moffett et al. 2025). Because interchromosomal duplicates are smaller, they are less likely to include the parent gene’s cis-regulatory elements, leaving the parent constrained and the daughter free to diverge; by contrast, larger intrachromosomal duplicates more often retain regulatory sequences, so both copies begin under similar constraint.

### Rapidly evolving segmental duplication landscapes

To quantify the segmental duplication landscapes, we constructed a separate segmental duplication network for each species, where nodes represent genomic regions and edges represent duplications. Edge weights indicate similarity scores, representing how recently the duplication emerged. For each species, we calculated 13 network properties and the average similarity score across duplications (a non-network property). We used these 14 measures collectively to quantify the segmental duplication landscape for that species (**see Methods**).

To assess whether our measures of segmental duplication landscapes were influenced by assembly qualities, we calculated correlations between each of the 14 landscape metrics and four assembly quality indicators: coverage width, contig NG50, scaffold NG50, and percentage of the assembly composed of unplaced contigs. Only four of the 14 measures (network density, weighted network density, 𝛽, and mean similarity score) were moderately correlated with at least one assembly quality metric (|𝑟| ≥ 0.3 and raw 𝑝 < 0.01). Thus, most features of the SD landscape are largely independent of assembly quality (**Figures S4-7**), indicating that our cross-species comparisons are robust.

Next, conducted comparative analyses of segmental duplication landscapes across the 118 genomes to investigate how genomic structural evolvability varies across vertebrate species. Specifically, we tested three hypotheses: (1) the “selective constraint hypothesis,” which states that segmental duplication patterns are highly conserved across species, showing minimal variation; (2) the “phylogenetic drift hypothesis,” which states that the segmental duplication landscape varies in accordance with phylogenetic distance between species; and (3) the “species-specific dynamics hypothesis” stating that the segmental duplication landscape evolves so rapidly that that it does not correlate with phylogeny (**Figure 4A**). We note that these three hypotheses are not mutually exclusive, but it is possible to parse which of these three is most dominant in shaping segmental duplication landscapes.

**Figure 4.**
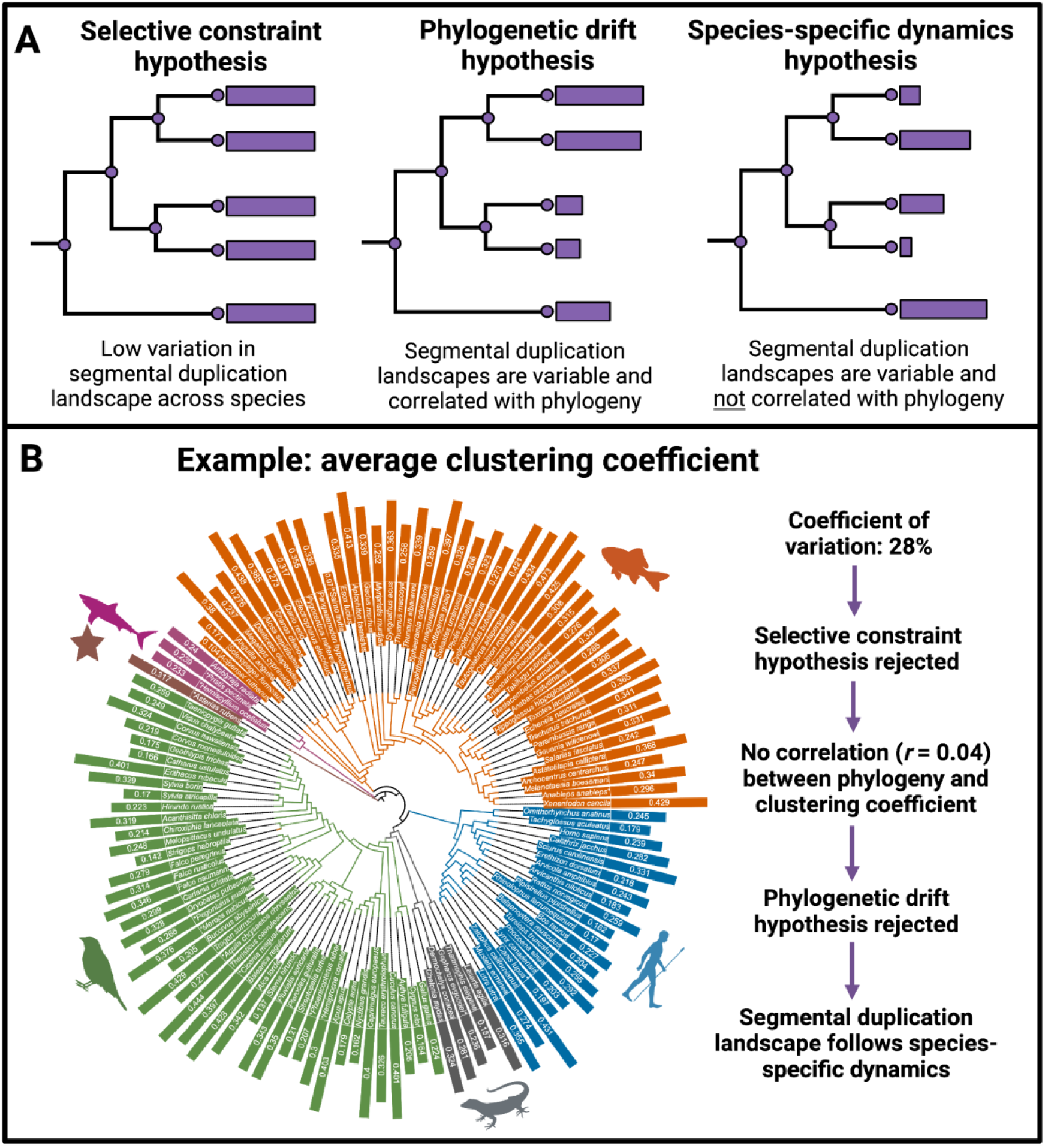
Hypotheses for the evolution of the segmental duplication landscape. **A.** Expectation for variation in the segmental duplication landscape under each hypothesis. The leaves in the trees in each schematic represent species. The length of the purple bar represents the value of a hypothetical measure of the segmental duplication for the corresponding species. The figure shows the expected variation in this measure under three hypotheses. **B.** A phylogenetic tree of 118 species annotated with the average clustering coefficient values. The height of the bar on each species indicates the value of the average clustering coefficient. Species’ names appended with an asterisk mean that a close substitute for the species was used to generate the tree (see **Table S2**). The variation in the average clustering coefficient across species matches the expectation under the species-specific dynamics hypothesis.

To test the selective constraint hypothesis (that evolutionary pressure conserves segmental duplication patterns across species), we analyzed all 14 measures across species. Following previous studies (Al-Sallami et al. 2014; Kumar et al. 2018), we considered a measure’s variation across species to be low if its coefficient of variation across species was less than 10%. We found that none of the segmental duplication landscape measures exhibited low variation (**Table S1**), suggesting that segmental duplication landscapes are not conserved in vertebrates. In fact, six out of the 14 measures, including the weighted density and mean node strength, exhibit coefficients of variation exceeding 100%. Based on these findings, we reject the selective constraint hypothesis and shift focus to the two alternative hypotheses (i.e., phylogenetic drift and species-specific dynamics) to explain the observed variation in segmental duplication landscapes.

To differentiate between the phylogenetic drift hypothesis and the species-specific dynamics hypothesis, we calculated pairwise distances for each of the 14 segmental duplication measures across species pairs. We correlated these with phylogenetic distances obtained from TimeTree (**see Methods**) (Kumar et al. 2022). If a species was not available in TimeTree, we used a closely related substitute (**Table S2**). If most of these 14 measures correlate with phylogeny, we favor the phylogenetic drift hypothesis; otherwise, we favor the species-specific dynamics hypothesis.

We found that none of the 14 measures of segmental duplication landscape correlates with phylogeny across vertebrates (|𝑟| > 0.3; See **Methods** for justification of the threshold; **Table S3**). Instead, the evolution of segmental duplication landscapes largely follows species-specific dynamics. This lack of correlation led us to reject the phylogenetic drift hypothesis in favor of the species-specific dynamics hypothesis. We demonstrate this reasoning using the average clustering coefficient, one of the 14 measures, in **Figure 4B**. We found that even very closely related species pairs (time since divergence < 50 million years) exhibit no greater similarities in segmental duplication trends than more distantly related species pairs (Cohen’s |𝑑| < 0.2 across measures; **Table S4**). These findings suggest that the evolution of segmental duplication patterns, driven by species-specific dynamics, occurs at such a rapid pace that measures of duplication landscapes may differ significantly even between closely related species (for example, **Figure S8**).

We note that one previous study (Abdullaev et al. 2021) reported that segmental duplication landscapes follow the phylogenetic drift hypothesis instead of the species-specific hypothesis supported by our results. However, their conclusion was based on only nine species and one network property (component size distribution). With our large dataset of 118 species, we found that the component size distribution also follows species-specific dynamics, consistent with the results based on our 14 measures **(see Supplementary Text S1)**. This result suggests that the results of (Abdullaev et al. 2021) may be limited in generalizability due to restricted species sampling and focus on a single network property.

Nevertheless, an interesting pattern stands out within mammals, but not across all species, in terms of the average similarity scores (**Figures 5A-B**). In particular, therian mammals (mammals with live birth) exhibit much higher average similarity scores than monotremes (platypus and echidna, the egg-laying mammals) (Hedge’s 𝑔 = 6.45, 𝑝 = 0.01; **Figure 5C**), leading to a strong correlation between phylogeny and average similarity scores within mammals 𝑟 = 0.85; 𝑝 = 2.26 × 10^−53^; **see Methods**). Because higher similarity scores generally indicate more recent duplications, this result suggests that monotreme genomes may contain older duplications and fewer recent duplications than therian genomes. However, we can break down the dissimilarity score (1 – similarity score) into two parts: mismatch error, which reflects the number of substitutions between duplicates, and gap error, caused by insertions or deletions. Our results show that the gap score differs significantly between monotremes and therian mammals (Hedges 𝑔 = 6.45; Mann-Whitney U test, 𝑝 = 0.01), while average mismatch scores are similar across both groups (Mann-Whitney U test, 𝑝 = 0.38) **(Figures 5D-E)**. If monotreme duplications were indeed older, we would expect both their mismatch and gap scores to be higher. In contrast, the fact that only gap scores are elevated in monotremes suggests their genomes are more prone to insertions and deletions rather than simply containing older duplications. Two hypotheses could be tested to explain the higher gap score in monotremes: 1) larger density of processed pseudogenes in monotremes; and 2) a higher rate of indels in monotremes. Parsing this out would be an interesting topic for future research.

**Figure 5.**
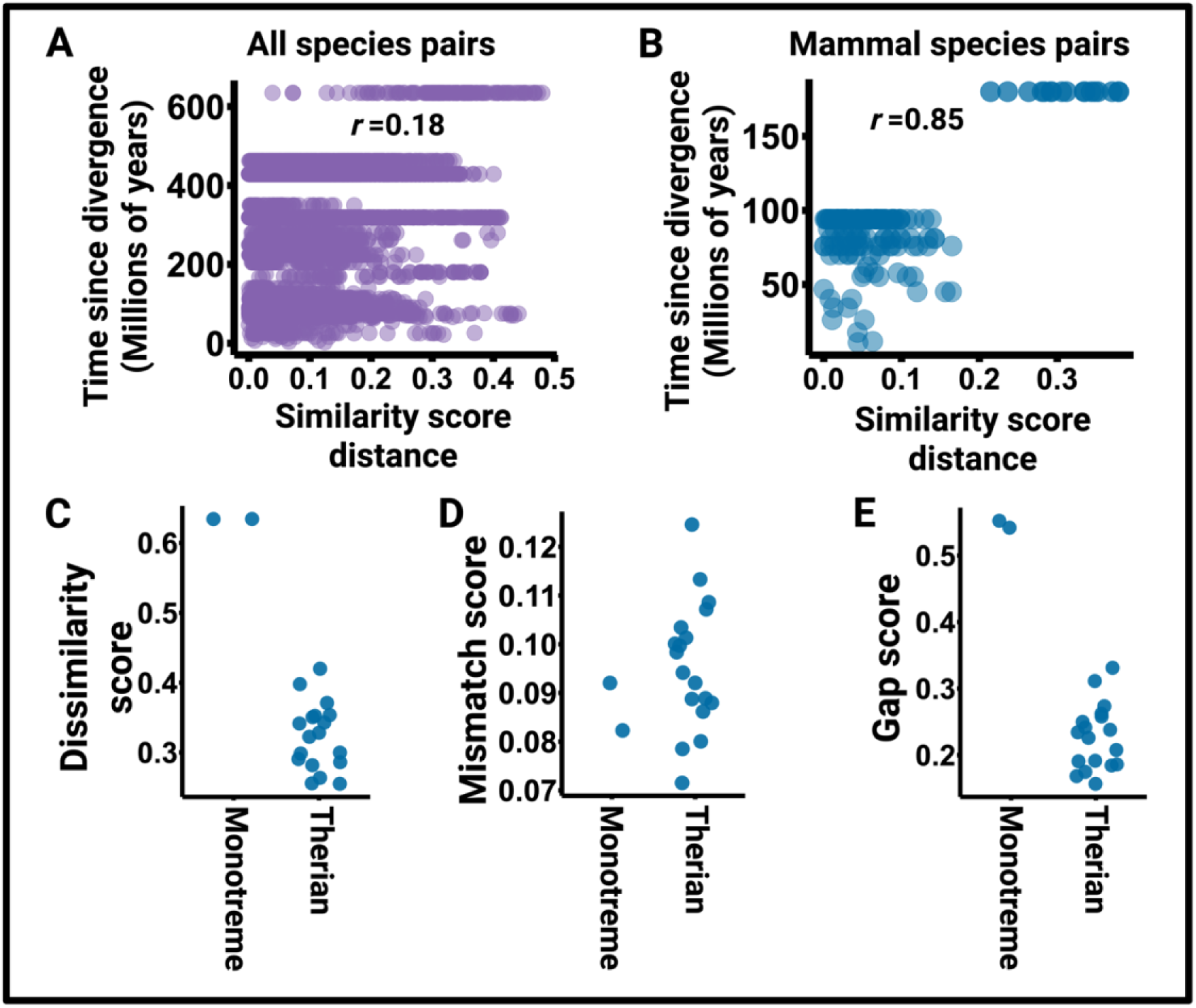
Insights from the average similarity scores for segmental duplications across species. **A.** Phylogenetic distances are not correlated with the average similarity score distances across pairs of species. **B.** Phylogenetic distances are strongly correlated with average similarity score distances across pairs of mammalian species. Monotreme-therian species pairs drive this correlation. In **A** and **B**, each point represents a species pair. **C-E.** Monotremes (echidna and platypus) have higher average dissimilarity scores (1 – similarity score) than therian mammals. This difference in average dissimilarity scores between therian mammals and monotremes is not driven by differences between mismatch scores but by differences in gap scores. In **C**-**E**, each point represents a mammalian species.

### Functional enrichments

We conducted Gene Ontology (GO) enrichment analyses using g:Profiler for 11 species represented in both our dataset and g:Profiler annotations. These 11 species included four mammals (human, platypus, dolphin, and greater horseshoe bat), four ray-finned fishes (eastern happy, brown trout, lumpfish, and climbing perch), two birds (kakapo and golden eagle), and one reptile (Good’s thornscrub tortoise). For each species, we identified the top 100 genes with the largest number of overlapping segmental duplications and tested these sets for enrichment of biological processes.

We did not detect any biological processes enriched across multiple species, suggesting that the types of genes amplified by segmental duplication are largely lineage-specific. Nonetheless, several species showed lineage-specific enrichments, including response to pheromones (𝑝-value adjusted for multiple hypothesis testing, 𝑝_adj_= 1.1 × 10^−11^) and sensory perception of chemical stimulus (𝑝_adj_ = 2.7 × 10^−8^) in platypus, regulation of DNA-templated transcription (𝑝_adj_ = 6.5 × 10^−11^) in humans, synaptic signaling (𝑝_adj_ = 0.0011) in greater horseshoe bat, protein metabolic process (𝑝_adj_ = 0.039) in dolphin, multicellular organismal process (𝑝_adj_ = 0.049) in brown trout, and regulation of lymphangiogenesis (𝑝_adj_ = 0.049) in lumpfish.

The case of the platypus is particularly striking: the enrichment is driven by the expansion of the vomeronasal pheromone receptor *V1R* gene family, consistent with previous findings (Grus et al. 2007; Zhou et al. 2021). Similarly, the functional enrichment in the greater horseshoe bat is driven by duplication of neural function-related genes, while in humans the functional enrichment of transcription regulation is driven by the expansion of zinc finger protein genes (**Figure 6**).

**Figure 6.**
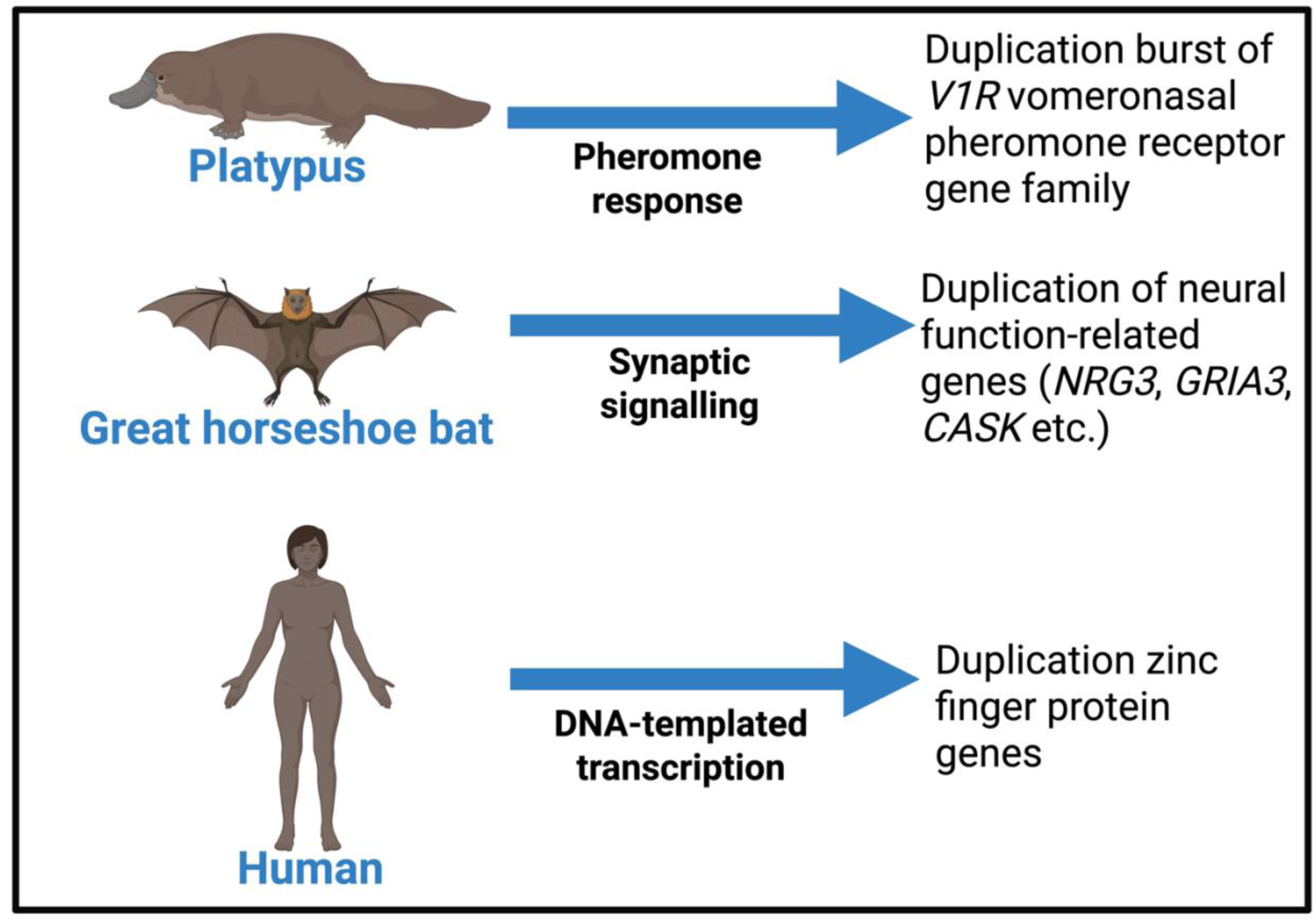
Functional enrichments of hyper-duplicated genes in the platypus, the great horseshoe bat, and humans.

## Discussion

In this study, we generate segmental duplication calls from repeat-masked long-read-sequenced genomes of 117 vertebrate species and the starfish. We caution that like most pipelines for detecting segmental duplications, our approach is sensitive to repeat masking. While we have used the most comprehensive repeat libraries available to mask genomes and indirectly show in our analysis that repeat masking is adequate, minor effects of inconsistencies in repeat masking across species may nonetheless linger.

Based on these segmental duplication calls, we find that vertebrate genomes exhibit a higher a propensity for tandem duplications relative to interspersed duplications, mirroring previous findings in humans. However, in subtelomeric regions, birds and mammals show the opposite propensity towards interspersed duplications. Moreover, we find that tandem duplications tend to be larger than interspersed duplications in vertebrates.

We quantified segmental duplication landscapes based on network measures extracted from segmental duplication networks. Comparing the differences in these properties against phylogenetic distances between species, we find that vertebrate segmental duplication landscapes are rapidly evolving. The rapid evolution of duplication landscapes across species may confer unique adaptive potentials, enabling certain lineages to respond more effectively to selective pressures. This may, in part, explain why some lineages are virtual living fossils while others are more prone to undergoing adaptive radiation.

Finally, we show that hyper-duplicated genes in certain species are enriched for specific biological processes. For example, in the platypus, these genes are enriched for pheromone response, driven by expansion of the vomeronasal pheromone receptor *V1R* gene family. Such expansions highlight how segmental duplications can become molecular substrates for evolutionary novelty.

A strength of our study is the application of network-based methods to study genomic structure. Network properties capture relationships among duplicated genomic regions that are invisible to traditional analyses, offering deeper systems-level insights into the structural organization of genomes.

Future work could extend our network framework to a multi-layer segmental duplication network, where each species represents a layer. Here, inter-species edges can connect orthologous genomic regions between species. This fine-grained approach, analogous to multi-layer co- expression networks (Russell et al. 2023), would enable the identification of communities of duplicated genomic regions that persist across layers (species). It would allow for the systematic discovery of gene copy number expansions underlying both lineage-specific adaptation and convergent evolution. A practical starting point could focus on a smaller and more closely related group of species such as primates, rodents, or carnivores, for which genomic alignments are more reliable. Overall, although analyzing multi-layer segmental duplication networks would be computationally demanding, it would provide deeper comparative insights than analyses limited to metrics derived from single-species segmental duplication networks.

As long-read sequencing technologies continue to improve and become commonplace, network-based approaches will become even more powerful tools for characterizing complex genomic phenomena. Overall, our study highlights the importance of integrating network analysis with evolutionary biology to unravel the intricate dynamics of genome evolution.

## Methods

### Data

We downloaded FASTA files for all 150 curated assemblies from GenomeArk (https://www.genomeark.org/), which holds genomes sequenced by the vertebrate genome project (as of August 16^th^, 2022). These assemblies included sex chromosomes. In this study, sex chromosomes and autosomes were analyzed together. For humans, we used the hg38 reference assembly for better comparison to previous studies on duplications in humans.

Before looking for segmental duplications (SDs), i.e., duplications with a size of at least one kilobase, it is necessary to mask all short repeats within each genome as much as possible to avoid false SD positives and improve the running time of SD detection software. This is because BISER (Išerić et al. 2022) (the SD detection software we use) assumes that all short repeats are filtered out; if they are not, short tandem repeat clusters and transposable elements will be called as SDs despite not being biologically so. Indeed, if these repeats are not masked, short repeats or transposable elements may masquerade as segmental duplications. We used RepeatMasker (v4.1.3) for this purpose. Since masking was inadequate for many non-mammalian species using the default masking library (Dfam minimal), we employed the complete Dfam 3.6 library. Additionally, we augmented this masking using species-specific repeat libraries from RepBase (v10/26/2018). Indeed, for all mammals, all birds, all reptiles, half of the ray-finned species, and 2 out of 3 of cartilaginous fish species, we used species-specific libraries. For the starfish, one cartilaginous fish, and the other half of the ray-finned fishes, species-specific libraries were not available. So we used class-level repeat libraries in addition to the complete Dfam library to mask them. Thus, the repeat masking is not expected to be highly inconsistent across taxa. Masking took from a few hours to a few weeks (it was especially performance-intensive on *Actinopterygii* genomes), and masked on average 25% of each genome: from 2% (*Asteroidea*) to 82% (*Bal. musculus*). We note that at the time of our analysis (data access date August 16th, 2022), the starfish genome was the only invertebrate genome available in the GenomeArk database.

Once the genomes were masked, we used BISER v1.2 (Išerić et al. 2022) to find all SDs within them. BISER relies on the SD error model (Numanagic et al. 2018) that assumes that (1) sequence similarity of SDs is higher than 75% (in other words, the error is below 25%), (2) random point mutations account for 15% of the total error and that they follow Poisson error distribution and are independent of each other (Jain et al. 2020); and (3) large indels and block variants account for the remaining 10% of the total error. The higher error rate threshold enables BISER to detect older SDs that occurred before the primate split in evolutionary history (the standard cutoff error threshold of 10% (Bailey and Eichler 2006) is sufficient only for detecting primate SDs (Numanagic et al. 2018)).

BISER finds all SDs in a given genome using a *k*-mer-based sketch of Jaccard distance through MinHash (Broder 1997) and *k*-mer windowing (Jain et al. 2020) to approximate edit distance under the described SD error model. This model allows it to quickly calculate SDs in the whole genome and decompose them into core blocks (Jiang et al. 2007). However, like all SD detection tools, BISER is extremely sensitive to the repeat masking quality; in our runs, it took from 4 minutes (good quality masking) to more than 10 days (low-quality masking; median time: 10 minutes) to find all SDs in a given genome. Our pipeline ran successfully for 118 genomes. All analysis in this manuscript is based on those 118 genomes. 109 of the 118 assemblies are pseudo-haploid consensus-collapsed assemblies. The remaining 9 are true (maternal or paternal) haploid assemblies obtained from trio-binning.

### Evolutionary distances

To get a phylogenetic tree for the 118 species analyzed here **(Figure 1A)**, we used a list of Latin binomials corresponding to these species and used it as input for TimeTree 5 (Kumar et al. 2022). On the tree, 14 species were replaced by close substitutes (**Table S2**). For our analysis, we used these close substitutes as proxies for the original species to measure phylogenetic distances. When using substitutes, we still use the names of the original species and append them with an asterisk. The output from TimeTree was a tree file in the Newick format with branch lengths in terms of the number of years. We used the `cophenetic.phylo()` function from the “ape” library in R (Paradis and Schliep 2018) to obtain evolutionary distances (in years) between each pair of species. The tree was visualized using the interactive Tree Of Life (iTOL) (Letunic and Bork 2021).

### Network construction

We construct a weighted network for each species from the data described above. We construct the network by considering consecutive windows, sized Δ = 5000 base pairs (5 kbp) on chromosomes **(Figure 7A)**. We use each such window as a node and connect pairs of nodes by weighted edges. We determine weighted edges as follows. In general, the duplication size does not coincide with Δ or multiples of it. For instance, assume that a region *R*_*x*_ of length 1.5Δ bp is duplicated in another region *R*_*y*_ of length 2.75Δ bp and that the similarity score between them is 𝑞. We assume that the starting coordinate of the region *R*_*x*_ corresponds to the starting coordinate of region *R*_*y*_, and the same for the ending coordinate. Therefore, a window of size Δ in *R*_*x*_ corresponds to a subregion of *R*_*y*_ of size 2.75Δ × (1Δ/1.5Δ) = 1.83Δ bp. For expository purposes, we also assume that *R*_*x*_ is contained in two nodes, 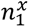 of size 1Δ bp and 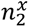 of size 0.5Δ bp, and that *R*_*y*_ is contained in three nodes, 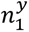 of size 0.85Δ bp, 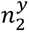 of size 1Δ bp, and 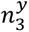 of size 0.9Δ bp **(Figure 7B)**. We calculate the edges, and their weights as follows. We assume that the size of *R_y_* is larger than *R*_*x*_ without loss of generality. First, we find the subregions of *R*_*y*_ that overlap the projection of the subregion of node 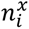 on *R*_*y*_, for each 𝑖th node contained in *R*_*x*_. This procedure determines the nodes in *R*_*y*_ that are adjacent to 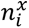 by an edge. For example, in **Figure 7B**, node 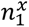 is projected to the first 1.83Δ bp of *R*_*y*_. Therefore, we connect 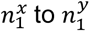 and 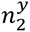 by an edge. Note that the entirety of 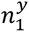 (of size 0.85Δ bp) overlaps with the projection of 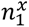 and that the first 0.98Δ bp of 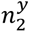 overlaps with the projection of 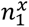. Likewise, 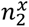 is projected to the last 0.92Δ bp of *R*_*y*_. Because 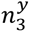 accounts for the last 0.9Δ bp of *R*_*y*_, we connect 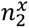 to each of 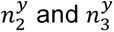 by an edge. Note that the last 0.02Δ bp of 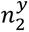 overlaps with the projection of 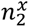 and that the entirety of 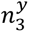 (of size 0.9Δ bp) overlaps with the projection of 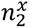. In this manner, we define an edge between a node in *R*_*x*_ and a node in *R*_*y*_ if and only if there is a positive overlap between *R*_*y*_ and the projection of *R*_*x*_ on *R*_*y*_. Second, we define the weight of the edge between 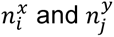 by

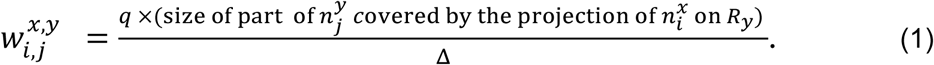

**Figure 7.**
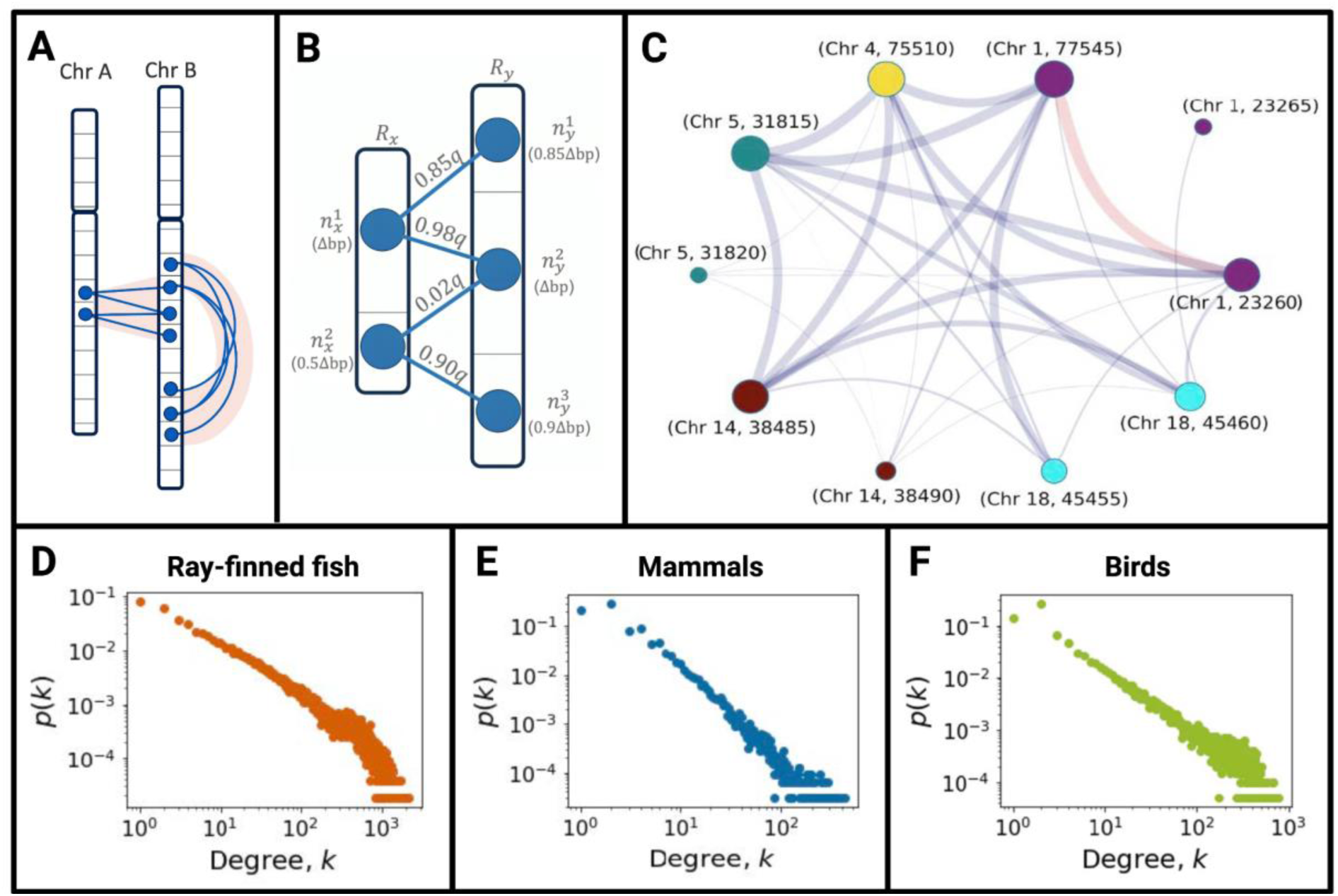
Network construction and degree distribution. **A.** Schematic for edge construction between two duplicated regions. **B.** Examples of edges for two chromosomes. The part of the network shown here is a magnification of edges between Chr A and Chr B in panel **A**. **C.** A part of the weighted segmental duplication network constructed for *Homo sapiens*. Different node colors represent different chromosomes. The number followed by the comma represents the starting coordinate (kbp) of the node on the chromosome **D-F.** The degree distribution of segmental duplication network from a representative species of ray-finned fish, mammals, and birds, respectively.

The generated networks are undirected. We allow a node to participate in edges with multiple duplications and edges with other nodes on the same or different chromosomes, as shown in **Figures 7A-B**. We show a part of the network of *Homo sapiens* in **Figure 7C**. The first component of each node label represents the chromosome number, and the second component represents the starting coordinate of the node in kbp. We recall that the node spans from the starting position to the subsequent 5 kbp. For example, node (Chr 1, 23260) is located on chromosome 1 and spans from 23260 kbp to 23264.999 kbp. We also show the degree distribution of three representative (chosen arbitrarily) networks from three taxonomic classes: ray-finned fish, mammals, and birds in **Figures 7D**, **E**, and **F**, respectively.

### Network properties

To understand the underlying duplication patterns, their evolutionary significance, and their potential impact on genomic architecture across species, we computed the following network properties for each species.

We denote the adjacency matrix of the segmental duplication network by 𝐴 = (𝑎_𝑖𝑗_), where the nodes are the genomic regions of size 5 kbp. The adjacency matrix represents whether each node pair is connected by an edge. We use the duplication similarity score as the weight of the edge between the 𝑖th and 𝑗th nodes, which we denote 𝑤_𝑖𝑗_. Note that 𝑎_𝑖𝑗_ = 𝑎_𝑗𝑖_ = 1 if 𝑤_𝑖𝑗_ = 𝑤_𝑗𝑖_ > 0, 𝑎_𝑖𝑗_ = 𝑎_𝑗𝑖_ = 0 if 𝑤_𝑖𝑗_ = 𝑤_𝑗𝑖_ = 0, and by assumption that 𝑎_𝑖𝑖_ = 𝑤_𝑖𝑖_ = 0. The degree of the 𝑖th node is defined by

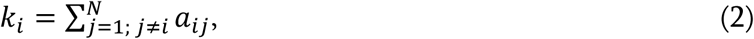

where 𝑁 is the number of nodes in the network. The degree represents the number of other nodes that the focal node is adjacent to by an edge. A high-degree node is a region that has been duplicated many times and, therefore, is suggested to be a hotspot of duplication activity.

The weighted degree, also called the node strength (Barrat et al. 2004), of the 𝑖th node is given by

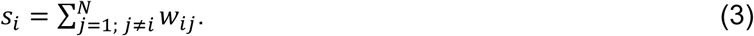

The node strength measures the cumulative similarity of all duplications for a focal genomic region across the genome. A node (region) with high strength is a region that has been duplicated many times and, on average, has high sequence similarity with its duplicates.

We have explained network metrics that are calculated for each node. We now explain other network metrics that are measured for each segmental duplication network, or equivalently, for each species. Some metrics use the node degree or the weighted degree of all the nodes in the segmental duplication network.

The mean degree is given by

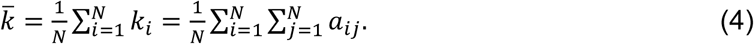

It is the average of the node degrees over all the nodes. It represents the average number of duplication events per genomic region. A higher mean degree indicates a more densely interconnected duplication network, possibly reflecting extensive segmental duplication events.

The mean node strength is given by

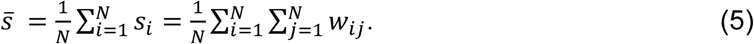

It is a measure of the duplication similarity score per pair of genomic regions. Larger 𝑠̅ values suggest stronger duplication relationships between genomic regions overall in the entire network.

We measure the heterogeneity of a distribution by the coefficient of variation (CV). The CV of the degree distribution and that of the strength distribution is given by

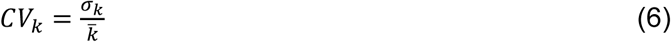

and

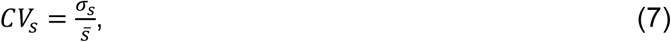

respectively, where 𝜎_𝑘_ and 𝜎_𝑠_ are the sample standard deviations of the node’s degree and strength, respectively. The CV of the degree distribution measures how much variation across the *N* nodes exists in terms of the number of duplication connections a node has. A large 𝐶𝑉_𝑘_ value indicates that some regions have markedly high degrees, representing duplication hubs, while other regions have small degrees. A small 𝐶𝑉_𝑘_ value suggests a relatively uniform duplication pattern across genomic regions. The CV of the node strength distribution measures the variation across the *N* nodes in terms of the cumulative duplication similarity score for a node. A large 𝐶𝑉_𝑠_ value suggests that some genomic regions have markedly strong duplication relationships with other regions. A small 𝐶𝑉_𝑠_ value indicates that duplication similarities are more evenly distributed across the genome.

The density of the network is defined by

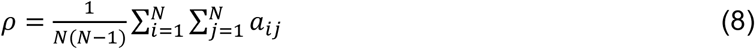

and represents the fraction of node pairs connected by an edge (Newman 2018; Wasserman 1994). High density means that segmental duplications are widespread, with many genomic region pairs sharing duplications. Low density suggests that duplications occur in a more isolated manner, possibly reflecting evolutionary constraints.

The weighted density of the network is given by

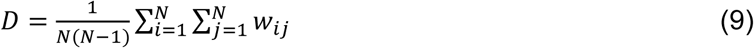

and represents the average edge weight over all node pairs, including those that are not adjacent by an edge (Wasserman 1994). The weighted density reflects the overall duplication similarity across the genome. A large 𝐷 value suggests that duplications tend to be strong in terms of the similarity score and widespread. A small 𝐷 value suggests that duplications are weaker or sparser.

The average clustering coefficient (Watts and Strogatz 1998) is defined by

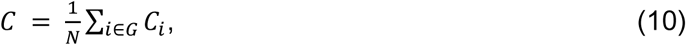

where 𝐶_𝑖_ is the local clustering coefficient for the 𝑖th node given by

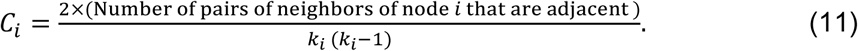

The average clustering coefficient represents the propensity that two neighbors of a node are also directly connected. A high *C* value suggests that duplications tend to form connected clusters in the sense of triangles, possibly indicating duplication hotspots. A low *C* value suggests that duplications occur in a more random fashion.

In addition to these properties, we also measure the mean, CV, and skewness of the edge weight, 𝑤_𝑖𝑗_ ^(^Watts and Strogatz 1998^)^. The mean edge weight represents the average similarity score between duplicated regions per duplication event. A high mean edge weight suggests recent or highly conserved duplications. A low mean weight suggests that duplications are generally divergent, suggesting older duplication events or functional divergence. The CV of the edge weight measures variability in duplication similarity across the duplicated region pairs. A large value of the CV of the edge weight implies a mix of old and new duplications. A small value of the CV of the edge weight suggests that the duplication similarity is relatively uniform across duplicated region pairs, indicating consistent duplication mechanisms for different duplicated region pairs. The skewness for the edge weight is given by

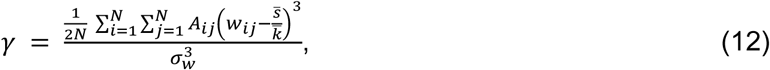

where 𝜎_𝑤_ is the sample standard deviation of the edge weight. The skewness of the edge weight measures the asymmetry in the distribution of the duplication similarity score (i.e., edge weight). Positive skewness (𝛾 > 0) suggests that most duplication events have low similarity, with a few having exceptionally high similarity. Negative skewness (𝛾 < 0) suggests that most duplications have high similarity, which may indicate genome regions with conserved segmental duplications. It should be noted that the CV also measures the extent of variation in the duplication similarity score; however, unlike the skewness, the CV does not measure the asymmetry.

We quantify the abundance and strength of cis-duplications relative to those of trans- duplications by measuring the so-called density ratio. To compute the density ratio for a given chromosome to other chromosomes, we first consider the density, 𝜌_cis_, of the subnetwork that consists of the edges that connect two nodes in the same chromosome. Second, we compute the density, 𝜌_trans_, of the subnetwork consisting of the edges connecting the node in the given chromosome to other chromosomes. Then, we compute the density ratio 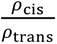 for the given chromosome. Finally, we calculate the mean of the density ratios over all chromosomes for a given species. We measure density ratios for both the unweighted network and the weighted network. A high unweighted density ratio implies that duplications mostly occur within the same chromosome. This case suggests localized duplication mechanisms such as tandem duplications or segmental duplications due to replication errors. A low unweighted density ratio implies that inter-chromosomal duplications are relatively frequent, which could be due to chromosomal translocations, rearrangements, or interspersed duplications. A high weighted density ratio signifies that intra-chromosomal duplications exhibit stronger similarity scores than inter-chromosomal ones overall. This case implies that intra-chromosomal duplications are either more conserved or more abundant than inter-chromosomal ones. A low weighted density ratio suggests that inter-chromosomal duplications have strong average similarity or are abundant.

We also explore how the node strength depends on the node degree for each network in a potentially nonlinear manner. We fit the following relation (Barrat et al. 2004):

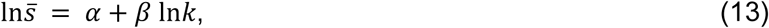

where 𝑘 is the node’s degree and 𝑠̅ is the node strength averaged over all the nodes whose degree is 𝑘. To estimate 𝛼 and 𝛽, we fitted a line to the set of points {ln𝑘, ln𝑠̅} using sklearn (Pedregosa et al. 2011) in Python. Equation (13) implies that 𝑠̅ is proportional to 𝑘^𝛽^. If 𝛽 > 1, then hubs, i.e., large-degree nodes, tend to have stronger duplications, implying that duplication hubs preferentially duplicate with highly similar regions. If 𝛽 = 1, there is no such correlation between the node’s degree and edge weight, implying that the duplication strength is independent of how many duplications a node has experienced. Various empirical weighted networks yield 𝛽 > 1 (Barrat et al. 2004).

To calculate the network properties, we use NetworkX version 2.8.5 in Python (Hagberg et al. 2008).

### Comparison with phylogenetic tree

We calculate all the 14 properties for each species in the dataset. For a given property, which is a scalar, we quantify the dissimilarity between two species 𝑝 and 𝑝′ in terms of the property by

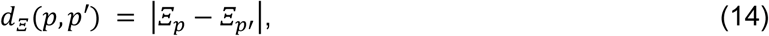

where 𝛯_𝑝_ and 𝛯_𝑝′_ represent the network property for species 𝑝 and 𝑝′, respectively. We compare 𝑑_𝛯_(𝑝, 𝑝′) to the phylogenetic distance between 𝑝 and 𝑝′ to explore which network-based measure may be associated with the phylogeny. We measure the correlation between the phylogenetic distance and 𝑑_𝛯_(𝑝, 𝑝′) by regarding each pair of species as a sample.

Here, we note an important concern. Our correlation analysis involves 6,962 species pairs (118 x 117/2). It is well-recognized in statistics that with such a large number of data points, hypothesis testing could yield “significant” results even for trivial effects (Berkson 1938; Cohen 1994). This phenomenon (“crud factor”) (Clayton 2021) cautions against overinterpreting 𝑝-values in large datasets. Therefore, rather than relying solely on 𝑝-value thresholds, we consider the two variables to be correlated only if |𝑟| > 0.3, the conventional threshold for medium effect size (Cohen 1988) that is routinely used in biology (Fredlund et al. 2012; Friedman and Alm 2012; Pai et al. 2019; Polk et al. 2025).

For the comparison in taxonomic class level, we omitted the analysis of the starfish, cartilaginous fish, and reptile classes because each of them had at most five species.

## Data Availability

The segmental duplication calls obtained using BISER, for the 118 genomes analyzed in this study have been deposited on FigShare. In addition to the files containing segmental duplication calls, the file “Species_and_Assembly_Info.csv,” containing taxonomic details of species, genome sizes, qualities of assembly, and the link to download the relevant FASTA (genomic sequence) files, is also available on FigShare. This data can be accessed using the following link:https://figshare.com/articles/dataset/Segmental_Duplication_Calls_BISER_Vertebrates_Lon <u>gReadGenomes/27859545</u>.

## Acknowledgment

I.N. was supported by the National Science and Engineering Council of Canada Discovery Grant (RGPIN-04973), Canada Research Chairs Program, Canada Foundation for Innovation’s John R. Evans Leaders Fund (CFI JELF) and B.C. Knowledge Development Fund (BCKDF). N.M. also acknowledges support from the Japan Science and Technology Agency (JST) Moonshot R&D (under grant no. JPMJMS2021), the National Science Foundation (under grant no.2052720), and JSPS KAKENHI (under grant nos.JP 23H03414, 24K14840, and W24K030130). O.G. acknowledges support from the National Science Foundation (under grant nos.2049947 and 2123284). O.G. and N.M. acknowledge support from the National Institute of General Medical Sciences (under grant no.1R01GM148973-01). The funders had no role in study design, data collection and analysis, publication decisions, or manuscript preparation.

## Supplementary Information

**Figure S1.**
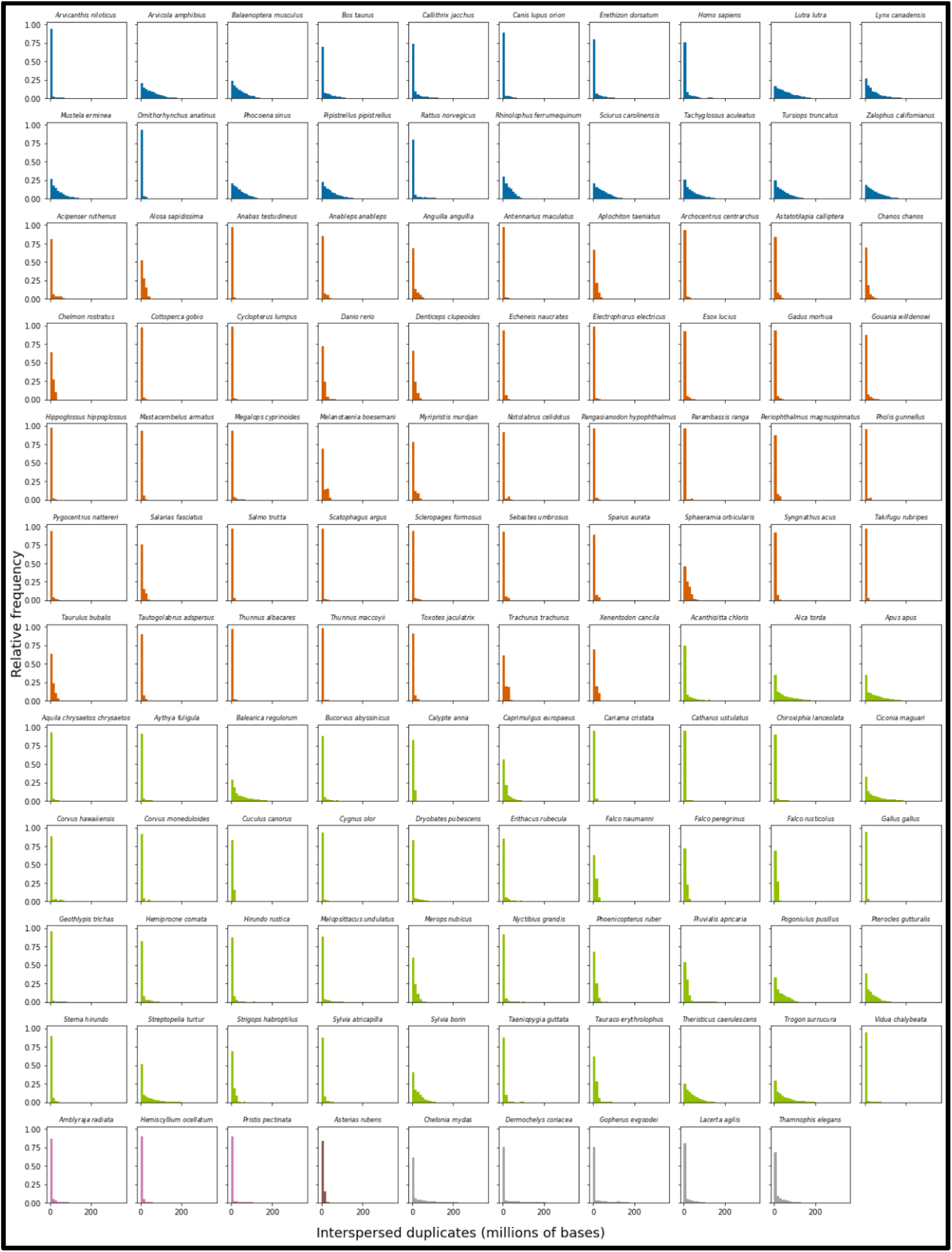
Distribution of distances between pairs of intrachromosomal duplicates. Each subplot represents distance distribution for one species. The histograms are color-coded by taxonomic class. The observed skewed distributions indicate that most duplicate pairs are located relatively close to each other, although some are separated by large genomic distances.

**Figure S2.**
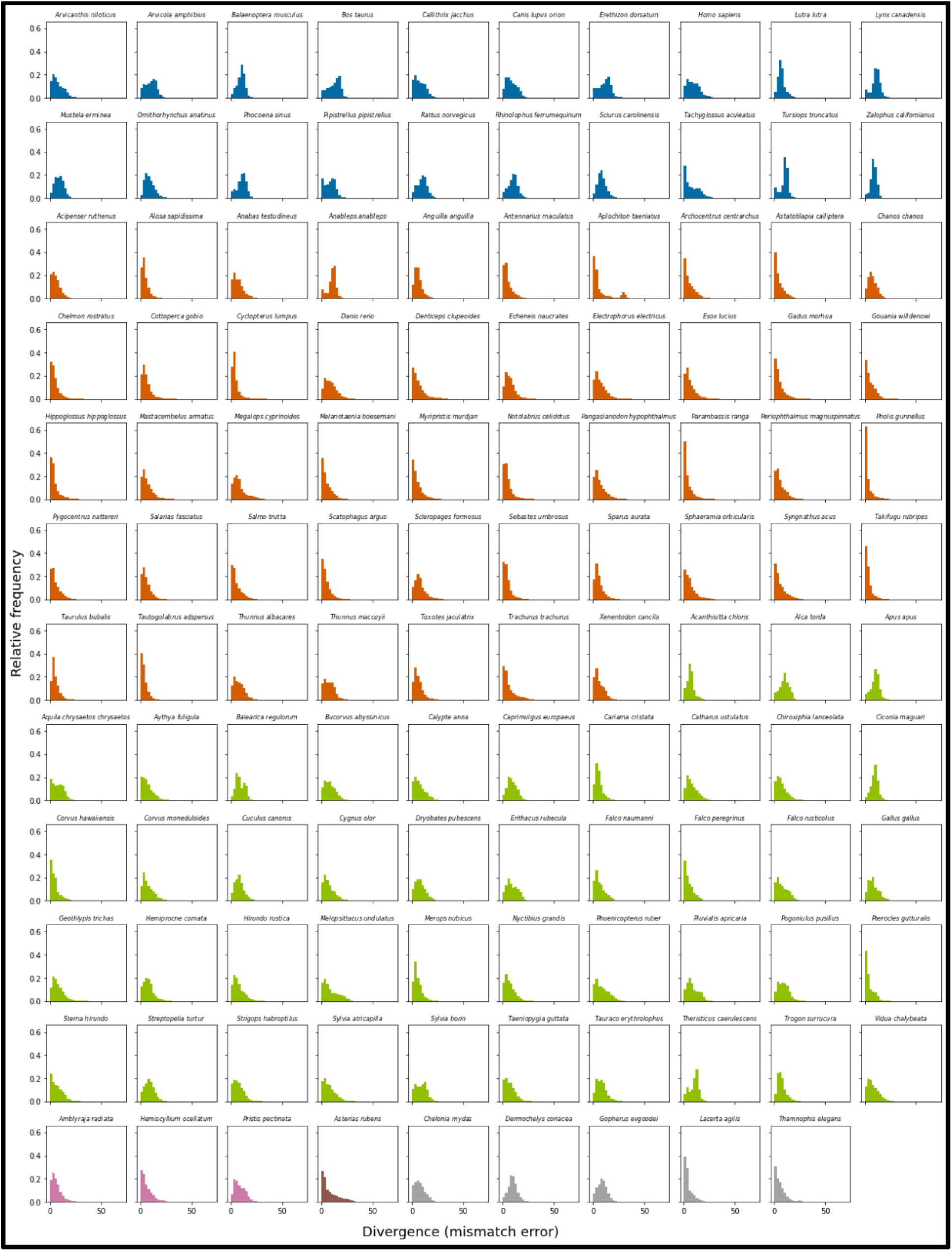
Divergence distributions of segmental duplications. We measure divergence as the mismatch score between partner duplicates. The histograms are color-coded by taxonomic class. We observe a general trend toward duplicates being more similar and younger. We note that this pattern is less pronounced for mammalian species than other species..

**Figure S3.**
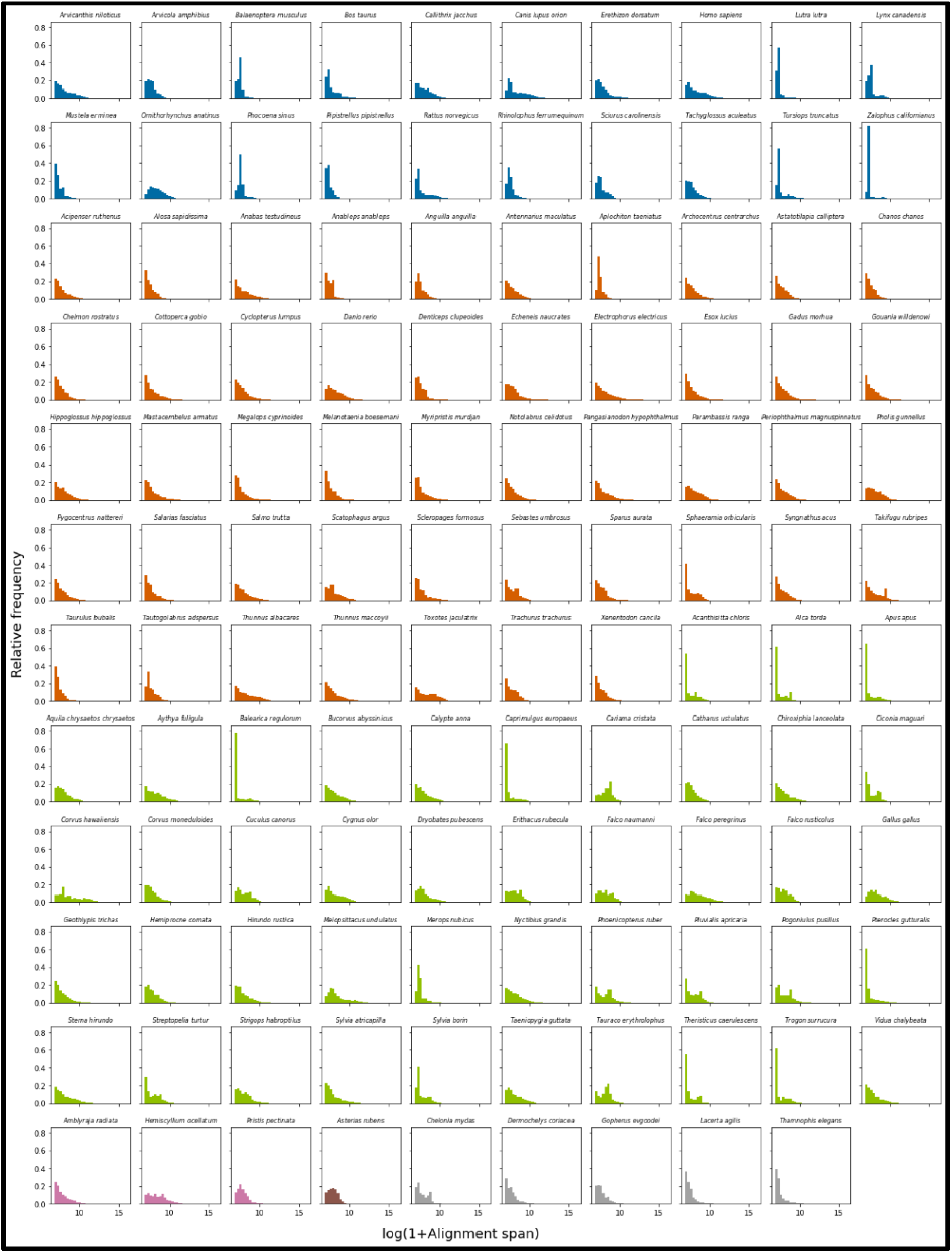
Size distributions of segmental duplications. The size (represented by the natural log of alignment span + 1) distribution of duplicate pairs for each species, color-coded by taxonomic classes. The observed distributions are typically right skewed, indicating a high proportion of shorter duplications with a long tail of large duplications, consistent across most species.

**Figure S4.**
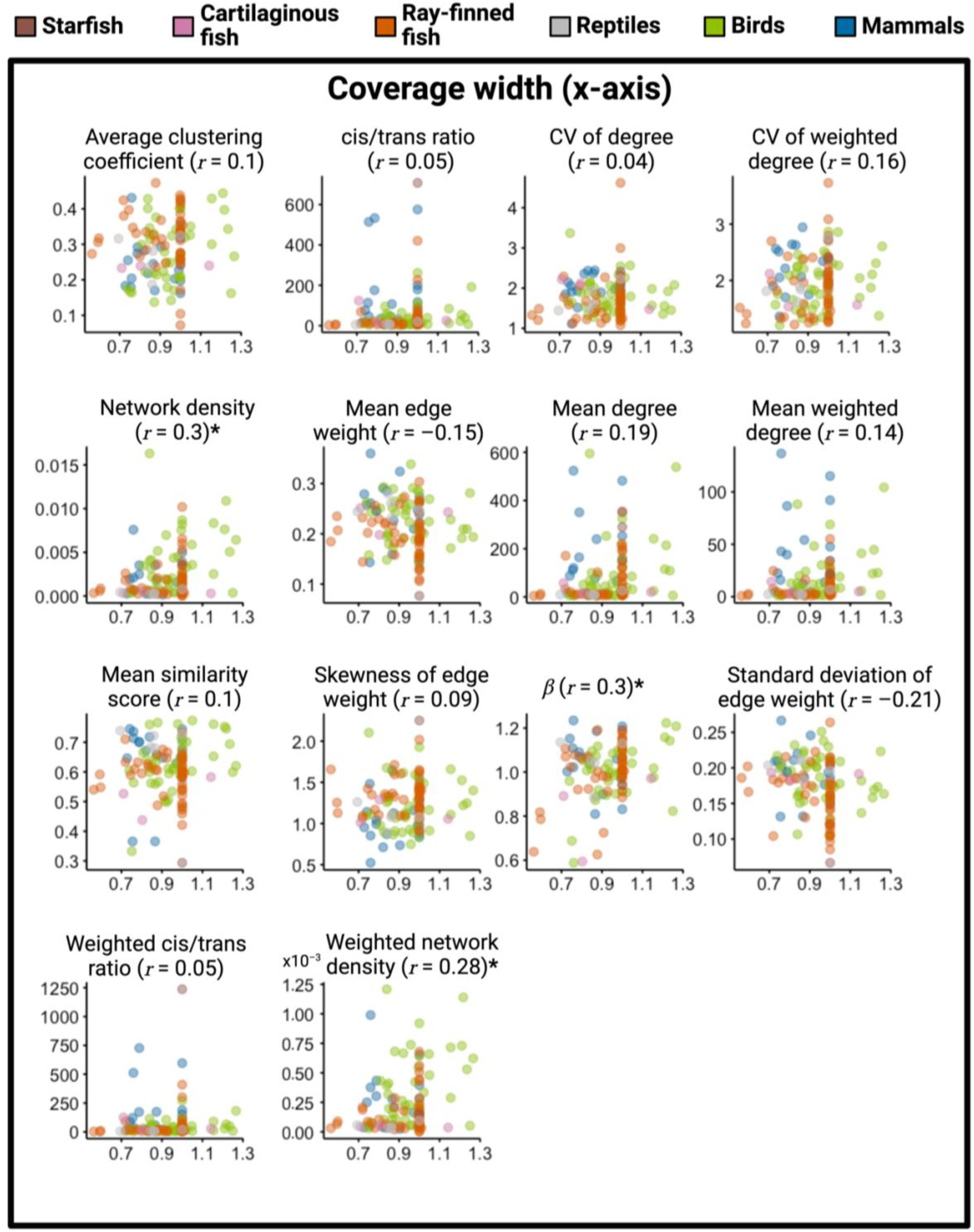
The relationship between coverage width and segmental duplication landscape metrics. The horizontal axis represents the coverage width. The vertical axis represents the segmental duplication landscape property whose name is shown on the top of each figure panel. A dot represents a species, color-coded by taxonomy class shown in the legend. Widths are greater than 1 when the assembled genome is larger than the estimated genome size. The Pearson correlation coefficient is denoted by 𝑟. Landscape properties for which the correlation has a raw 𝑝-value < 0.01 are appended by *.

**Figure S5.**
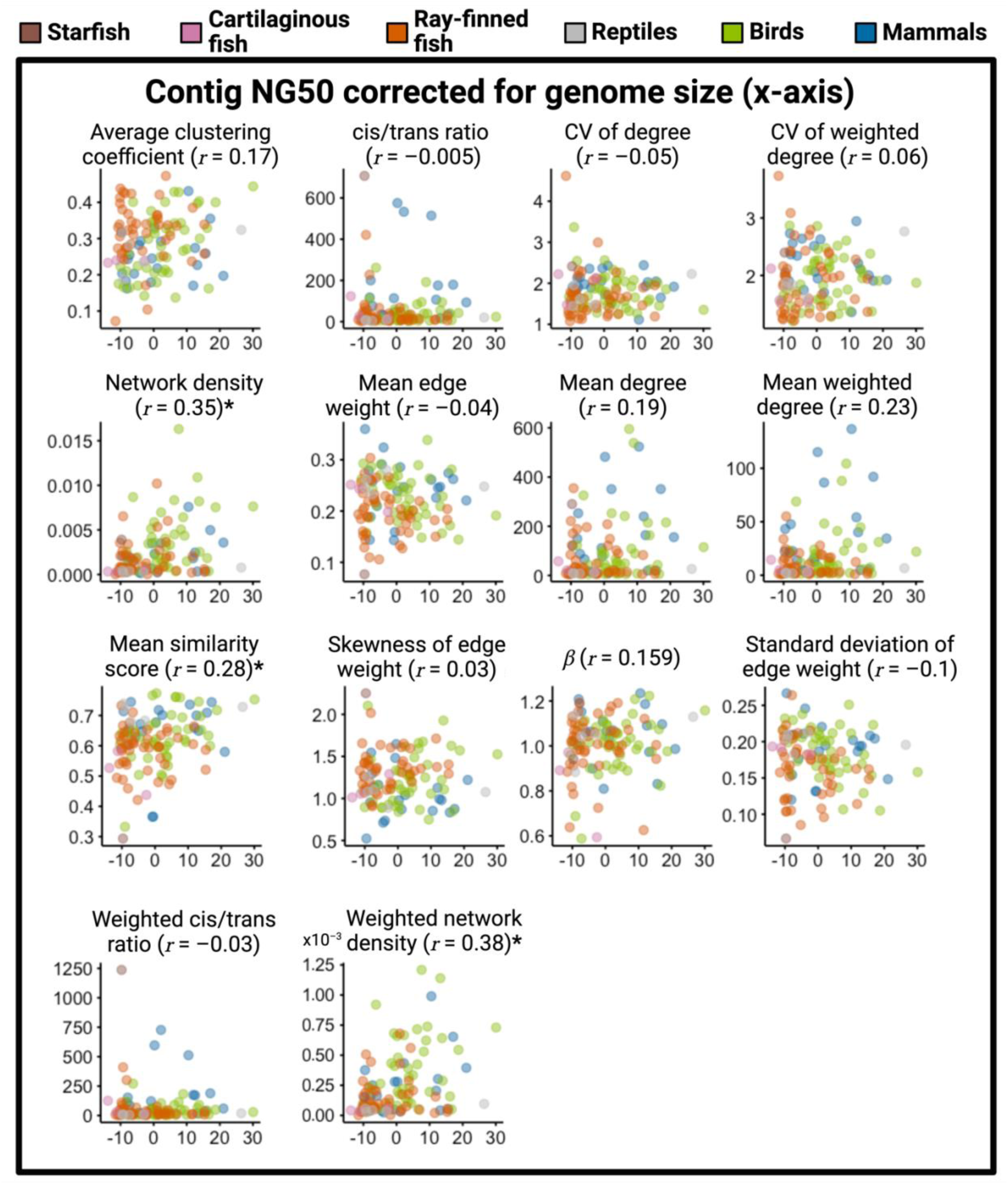
The relationship between contig NG50 corrected for genome size and segmental duplication landscape metrics. The horizontal axis represents the residuals from the linear model. The vertical axis represents the segmental duplication landscape property whose name is shown on the top of each figure panel. A dot represents a species, color-coded by taxonomy class shown in the legend. The Pearson correlation coefficient is denoted by 𝑟. Landscape properties for which the correlation has a raw 𝑝-value < 0.01 are appended by *.

**Figure S6.**
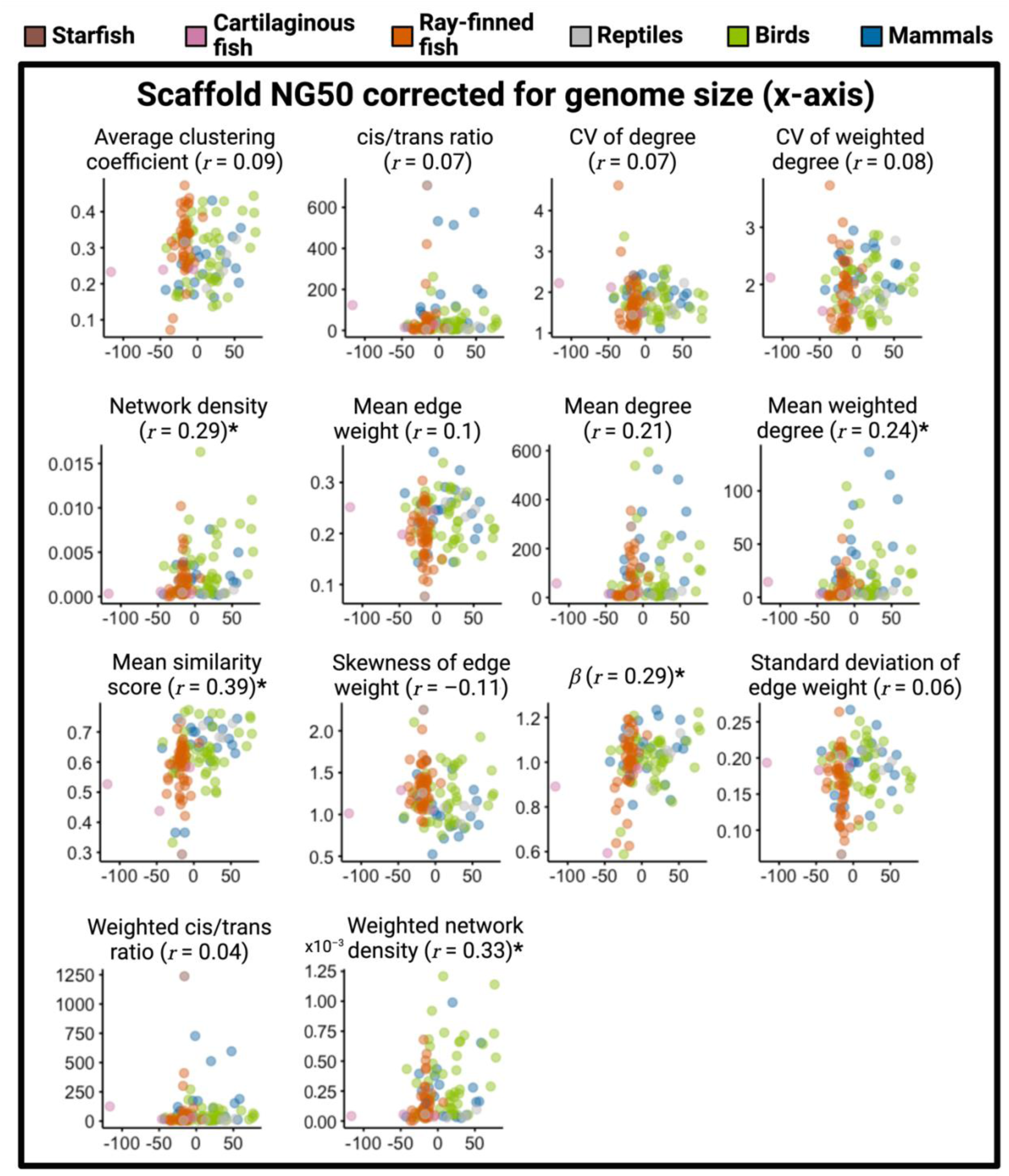
The relationship between scaffold NG50 corrected for genome size and segmental duplication landscape metrics. The horizontal axis represents the residuals from the linear model. The vertical axis represents the segmental duplication landscape property whose name is shown on the top of each figure panel. A dot represents a species, color-coded by taxonomy class shown in the legend. The Pearson correlation coefficient is denoted by 𝑟. Landscape properties for which the correlation has a raw 𝑝-value < 0.01 are appended by *.

**Figure S7.**
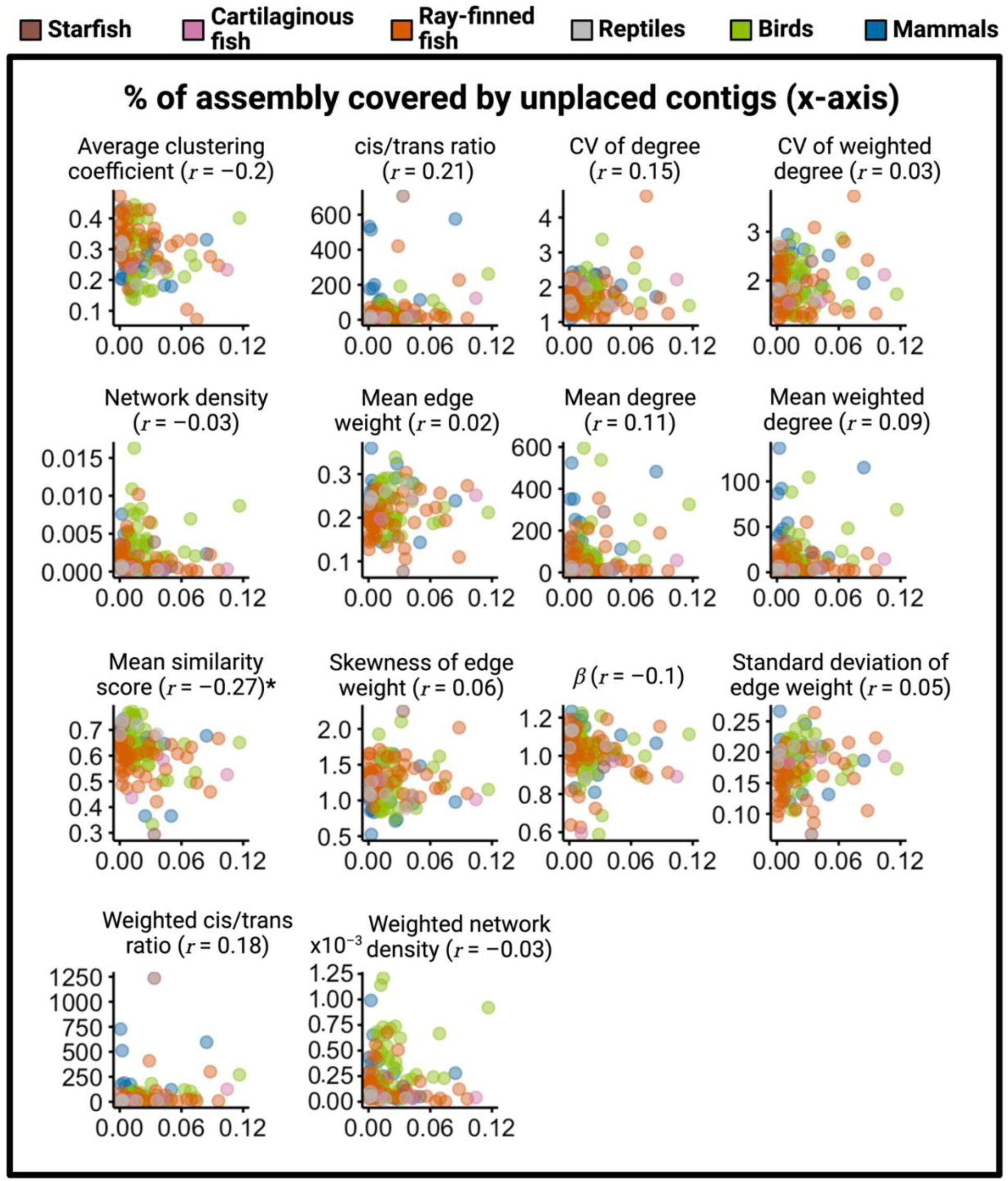
The relationship between the percentage of genome assembly covered by unplaced contigs and segmental duplication landscape metrics. The horizontal axis represents the unplaced contig coverage. The vertical axis represents the segmental duplication landscape property whose name is shown on the top of each figure panel. A dot represents a species, color-coded by taxonomy class shown in the legend. The Pearson correlation coefficient is denoted by 𝑟. Landscape properties for which the correlation has a raw 𝑝-value < 0.01 are appended by *.

**Figure S8.**
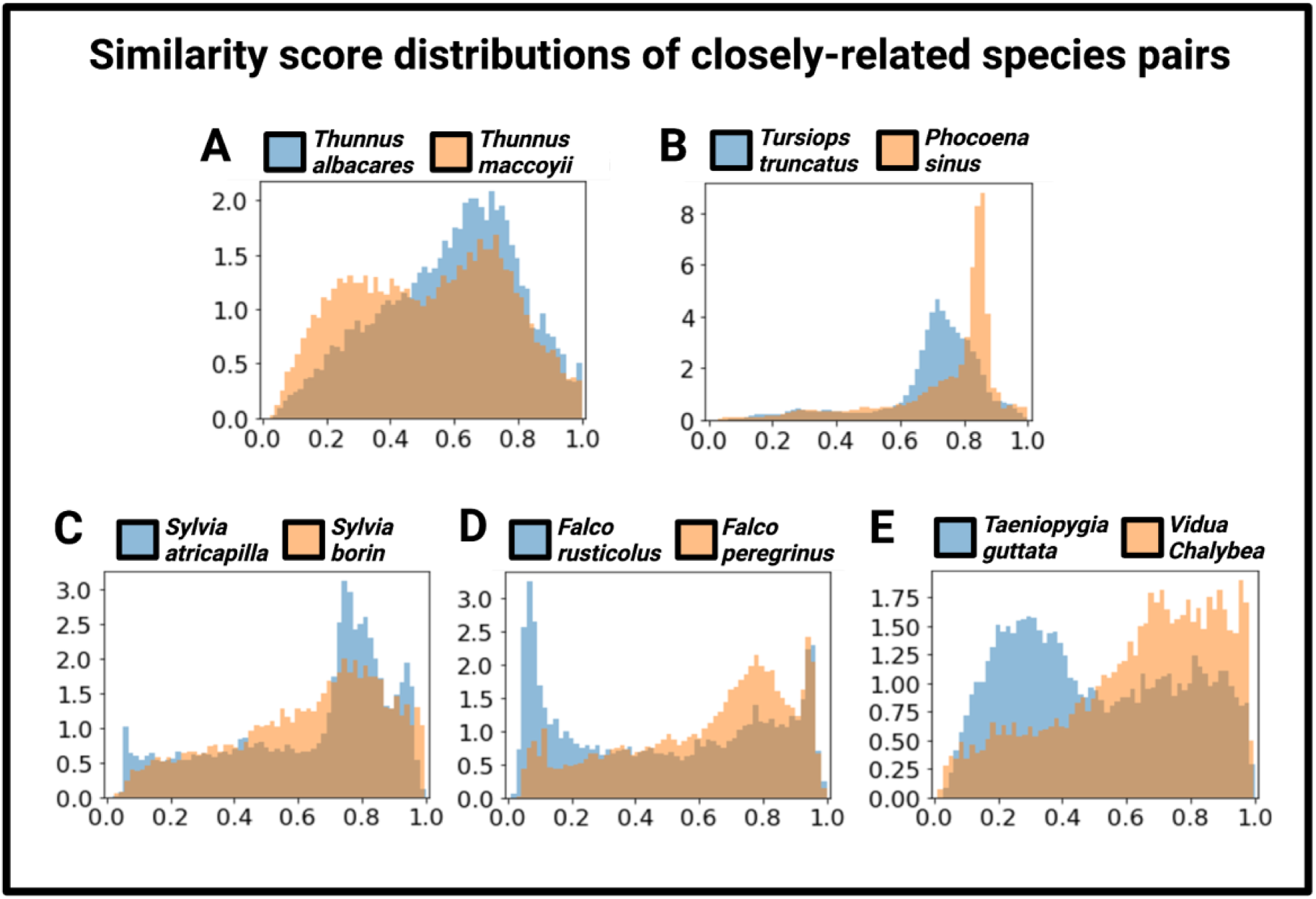
Similarity score distributions for closely-related species pairs. Five pairs of species that are phylogenetically close but have highly variable similarity distributions. The x-axis represents the similarity score and the y-axis represents density. **A.** Ray-finned fish. **B.** Mammals. **C-E.** Birds.

**Table S1.**
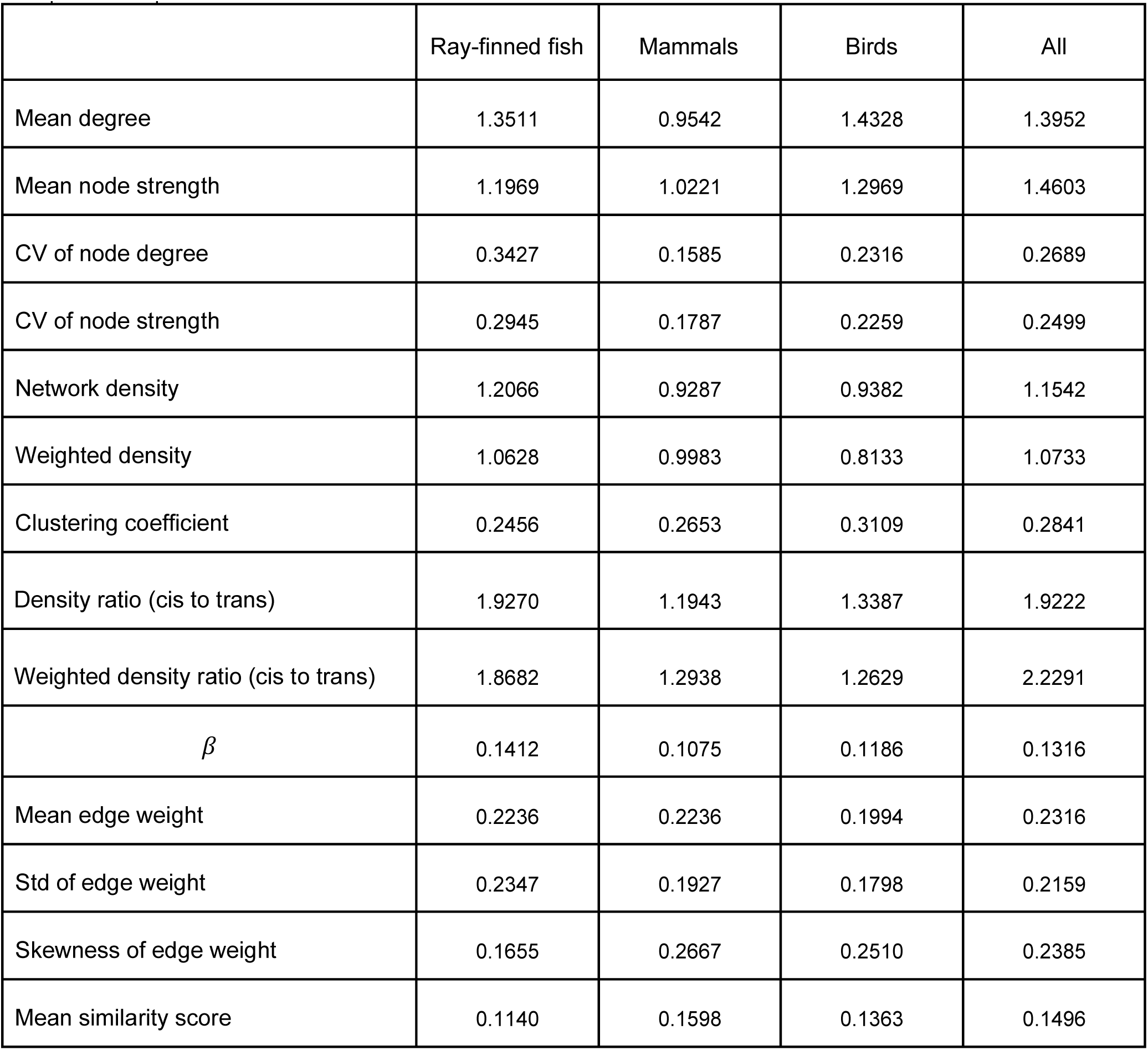
CV for each of the 14 measures across species. For example, the “CV of node degree’’, shown in the third row of the table, is a scalar measure calculated for each species. Its CV calculated based on all the ray-finned fish species is equal to 0.3427.

**Table S2.**
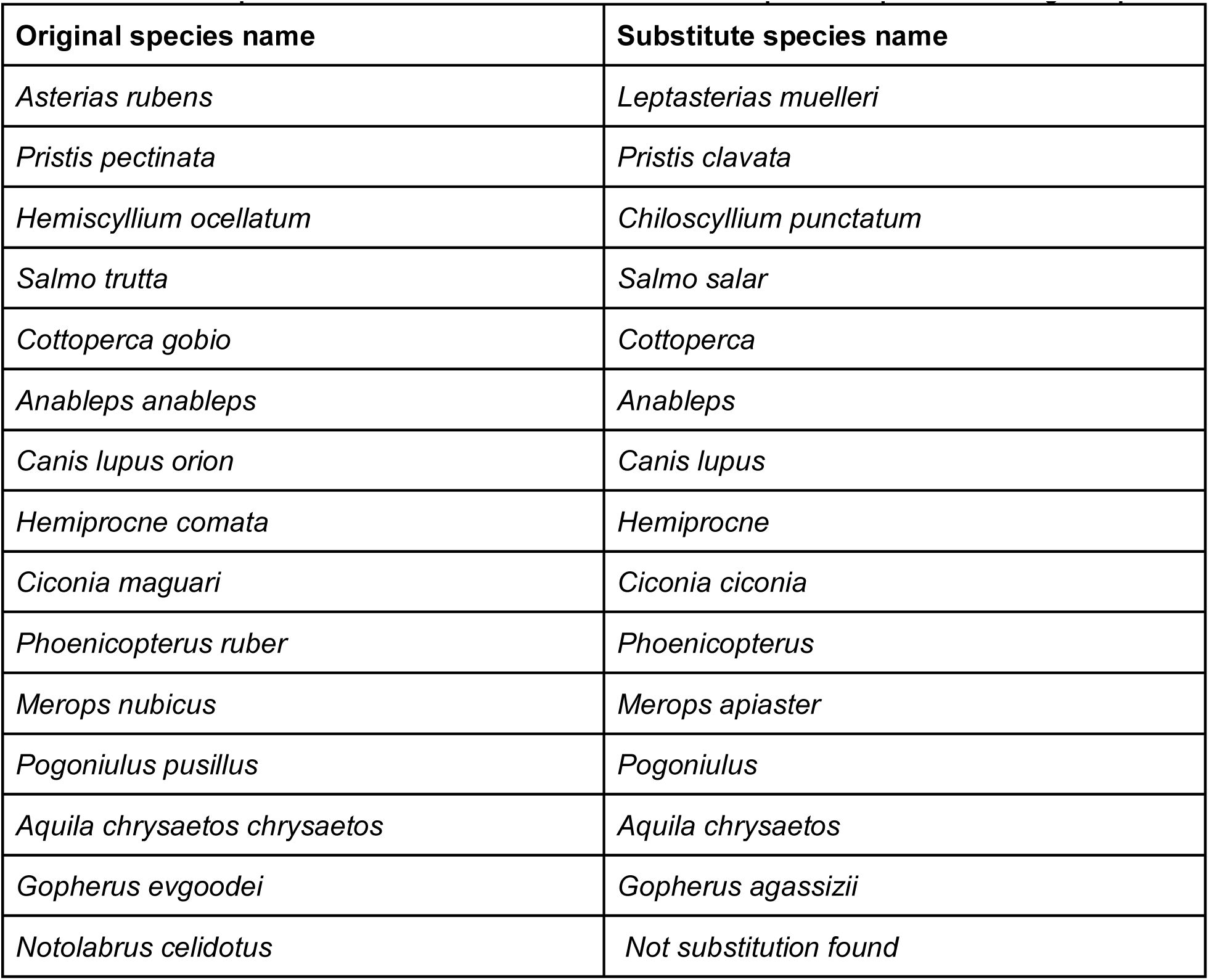
The list of species for which we use a close substitution as proxies in place of the original species.

**Table S3.**
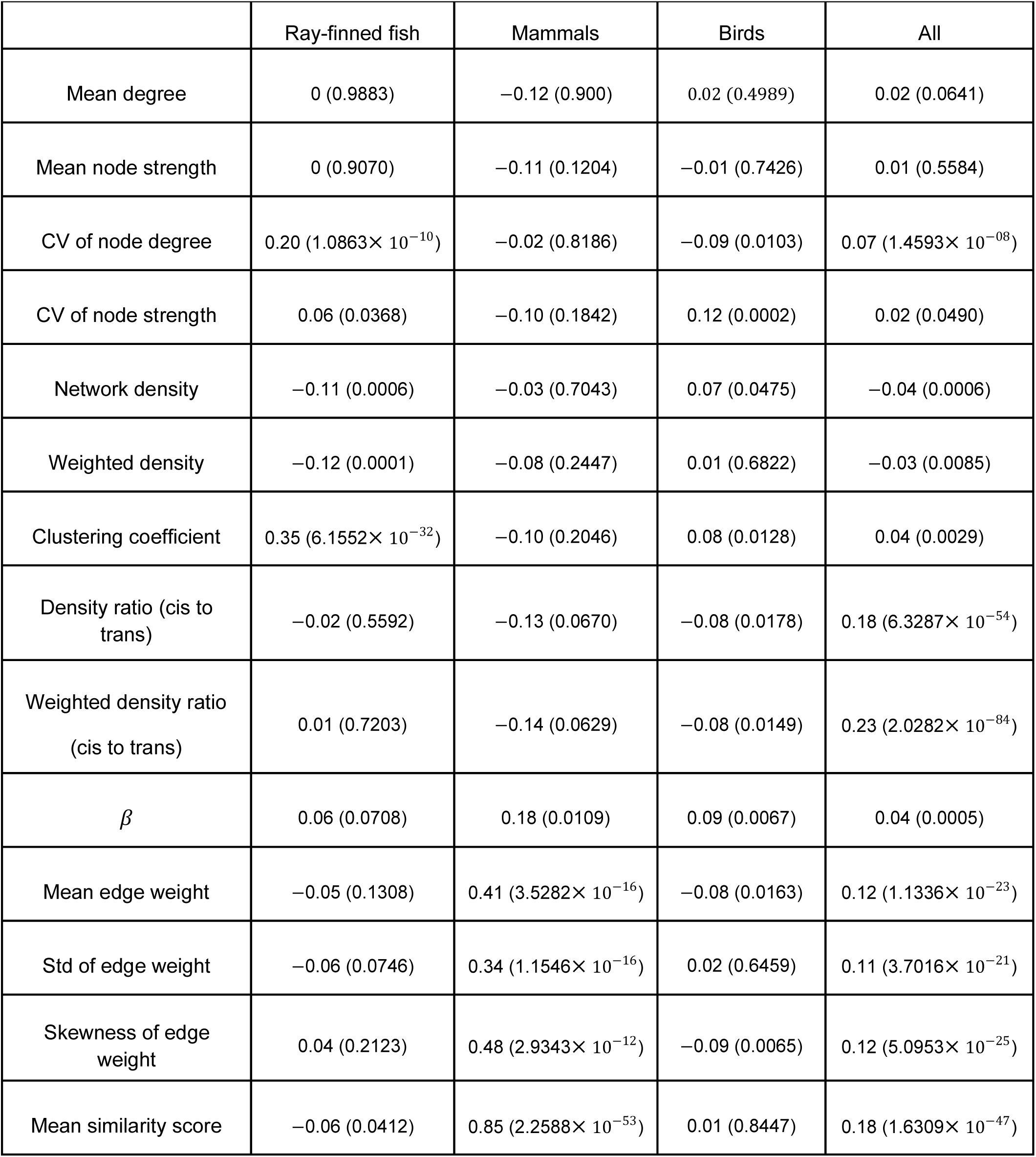
The correlation coefficient between the phylogenetic distance between two species and the distance in terms of each of the 14 measures. The corresponding 𝑝-value is shown in the parentheses.

**Table S4.**
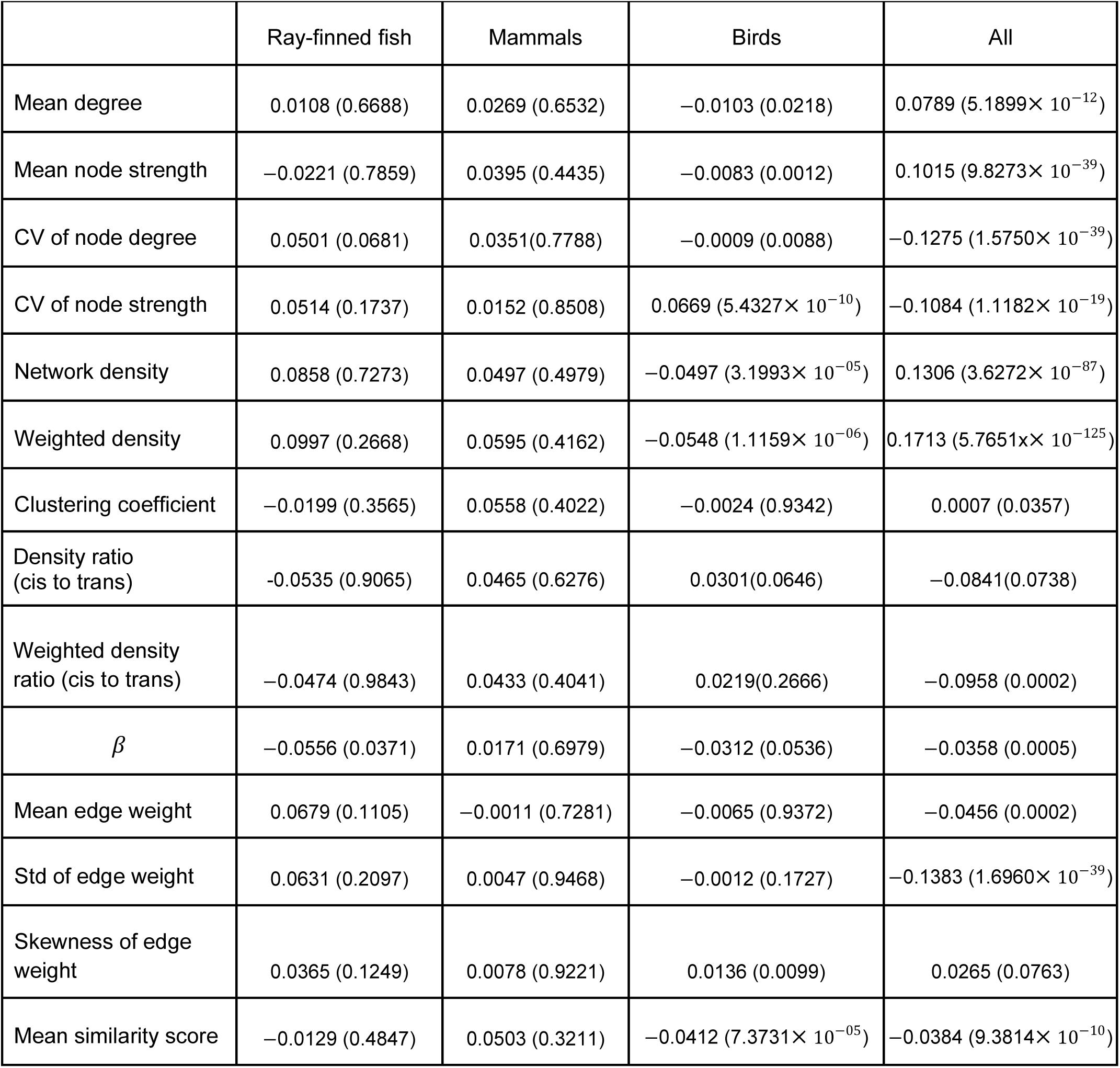
Effect sizes (Cohen’s 𝑑) and p-values from the Mann-Whitney U test comparing species pairs with phylogenetic divergence less than 100 million years to pairs with greater divergence.

## Supplementary Text S1

In (Abdullaev et al. 2021), the authors examined segmental duplication networks of nine species. They found that the component size distribution of the network was more similar within the six mammal species than between the mammal and non-mammal species. We computed the property that they used (i.e., the ordered sequence of the component size of the segmental duplication network) for each of our 118 species and computed the so-called Bray-Curtis dissimilarity (BCD) between each pair of species using the same method as theirs (Abdullaev et al. 2021). Then, as we did for our 14 quantities in the main analysis, we calculated the Pearson correlation coefficient between the phylogenetic distance between pairs of species and the BCD. The correlation was 𝑟 = –0.18 (𝑝 = 5.1905 × 10^−09^), 𝑟 = 0.23 (𝑝 = 0.0015), 𝑟 = 0.05 (𝑝 = 0.1635), and 𝑟 = 0.14 (𝑝 = 4.5435 × 10^−33^) for ray-finned fish, mammals, birds, and all species, respectively. Although the *p* values are mostly small due to large sample sizes, 𝑟 (as a measure of the effect size) is small. This result contrasts with those in (Abdullaev et al. 2021), which suggested that the component size distribution has an evolutionary signal, i.e., it is more conserved within mammals than between mammals and non-mammals. There are at least two possible reasons for the difference between their results and ours. First, we mainly compared within each of the three clades and used as many as 118 species. In contrast, their mammal versus non-mammal comparison is only based on six mammal species and three non-mammal species. Furthermore, one of the three non-mammal species used was *Caenorhabditis elegans*, which is evolutionarily farther than among the three clades we used. Second, the two studies used different computational methods to construct segmental duplication networks. Across-species comparison of segmental duplication networks aiming to improve the accuracy of segmental duplication calls and explicitly considering gene orthologs warrant future work.

