## Supplementary figures, tables, and text. for "Genomes from 117 vertebrate species reveal rapidly evolving segmental duplication landscapes"

### <u>Supplementary Information</u>

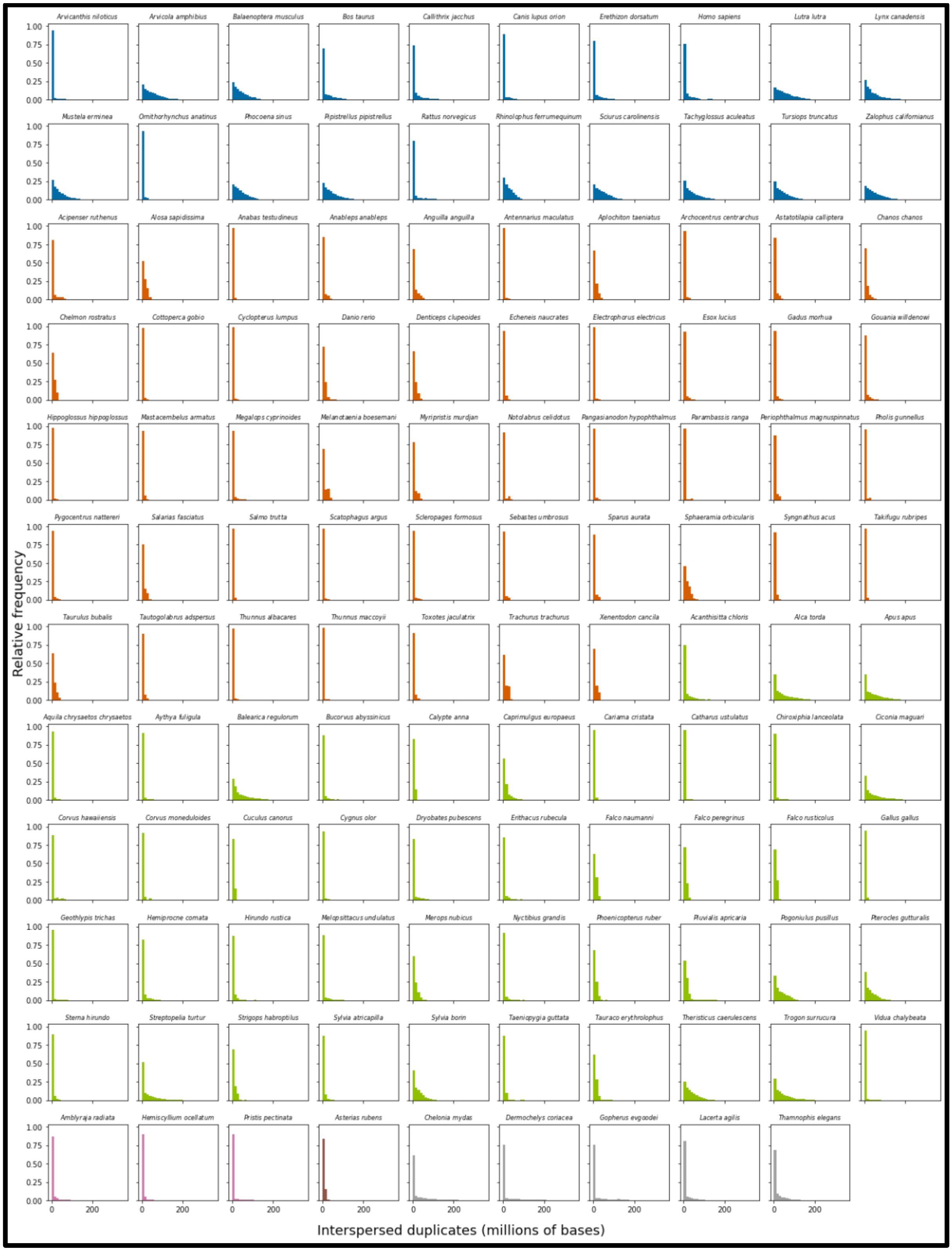


**Figure S1. Distribution of distances between pairs of intrachromosomal duplicates.** Each subplot represents distance distribution for one species. The histograms are color-coded by taxonomic class. The observed skewed distributions indicate that most duplicate pairs are located relatively close to each other, although some are separated by large genomic distances.

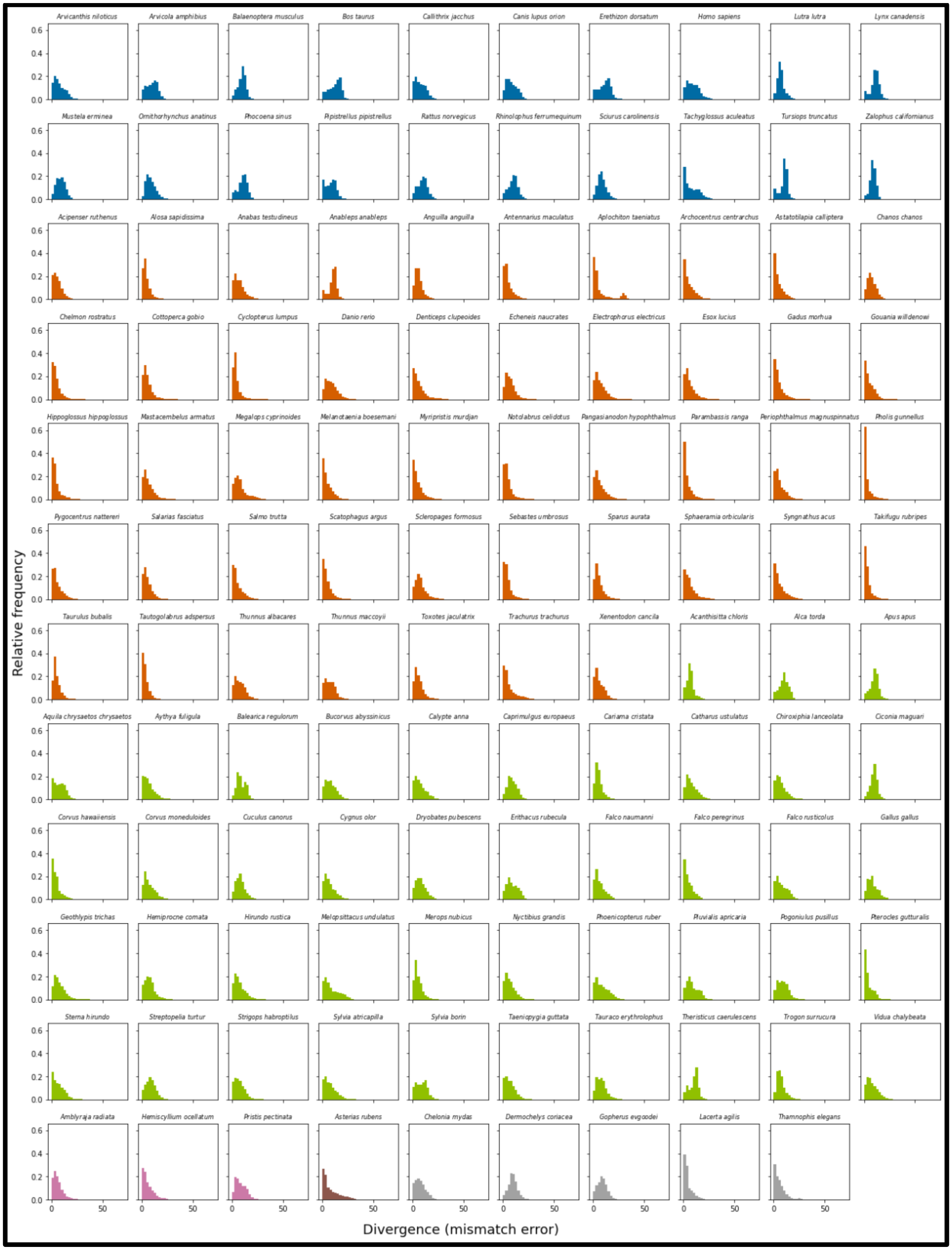


**Figure S2. Divergence distributions of segmental duplications.** We measure divergence as the mismatch score between partner duplicates. The histograms are color-coded by taxonomic class. We observe a general trend toward duplicates being more similar and younger. We note that this pattern is less pronounced for mammalian species than other species..

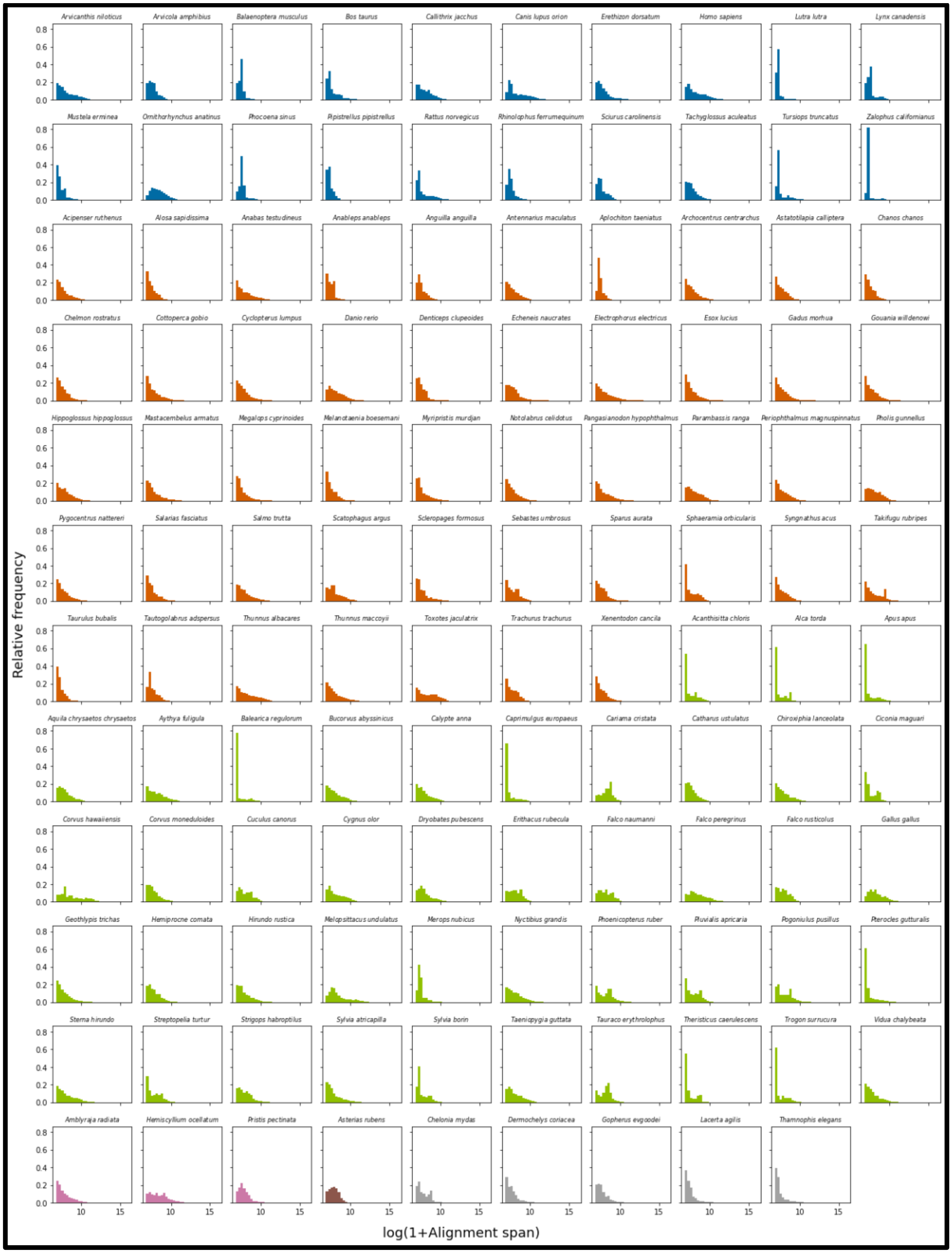


**Figure S3. Size distributions of segmental duplications.** The size (represented by the natural log of alignment span + 1) distribution of duplicate pairs for each species, color-coded by taxonomic classes. The observed distributions are typically right skewed, indicating a high proportion of shorter duplications with a long tail of large duplications, consistent across most species.

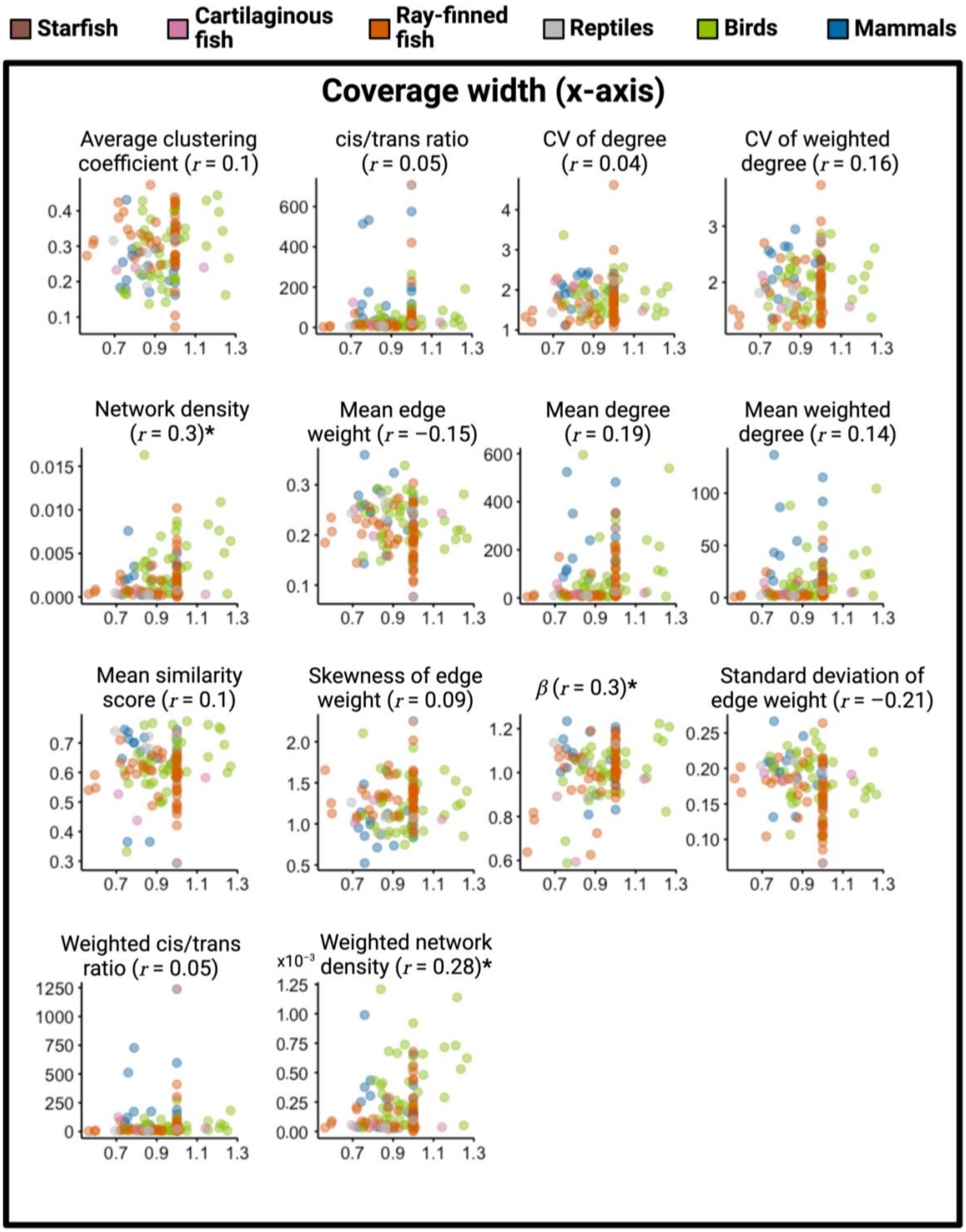


**Figure S4. The relationship between coverage width and segmental duplication landscape metrics**. The horizontal axis represents the coverage width. The vertical axis represents the segmental duplication landscape property whose name is shown on the top of each figure panel. A dot represents a species, color-coded by taxonomy class shown in the legend. Widths are greater than 1 when the assembled genome is larger than the estimated genome size. The Pearson correlation coefficient is denoted by $r$. Landscape properties for which the correlation has a raw $p$-value < 0.01 are appended by *.

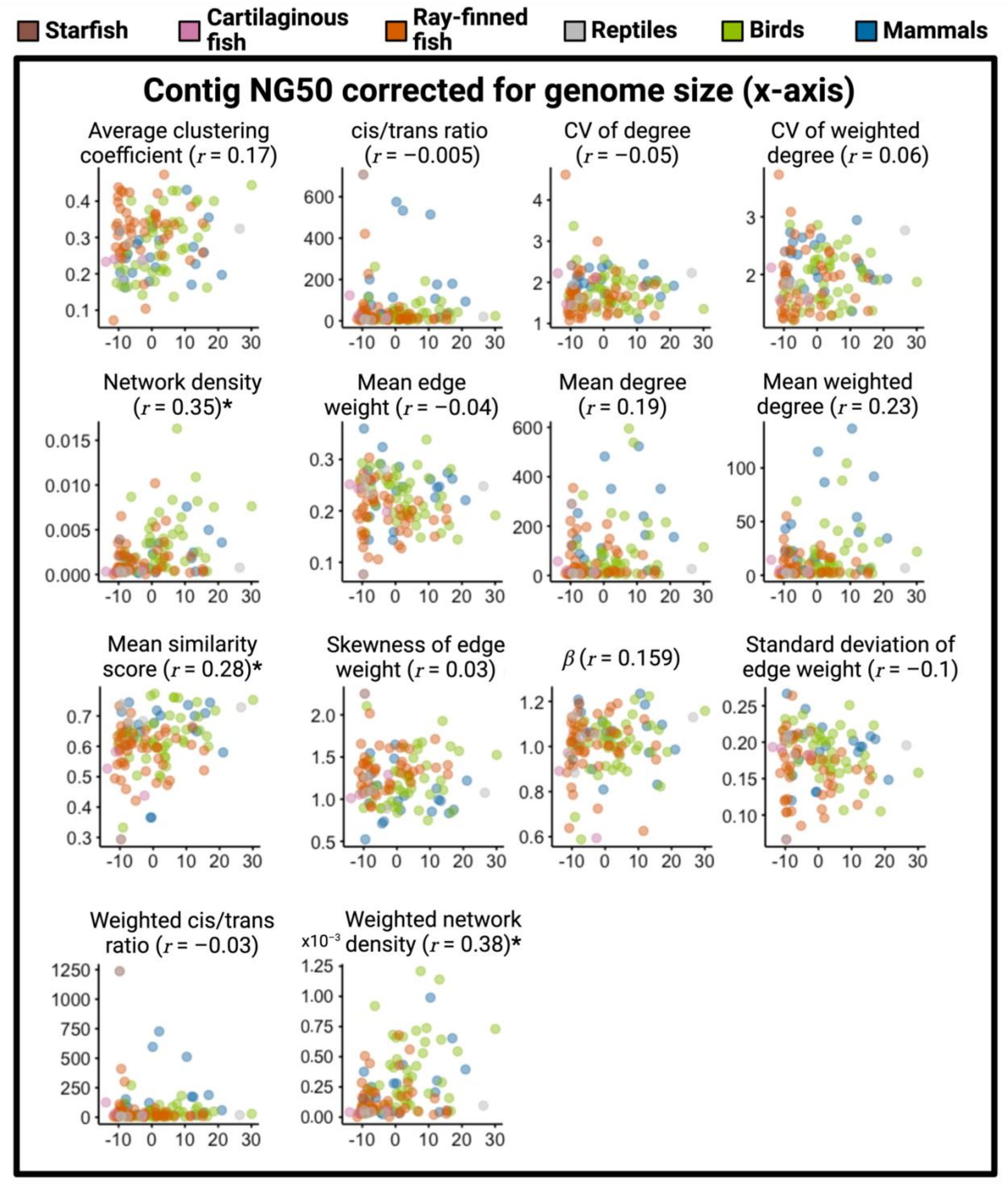


**Figure S5. The relationship between contig NG50 corrected for genome size and segmental duplication landscape metrics.** The horizontal axis represents the residuals from the linear model. The vertical axis represents the segmental duplication landscape property whose name is shown on the top of each figure panel. A dot represents a species, color-coded by taxonomy class shown in the legend. The Pearson correlation coefficient is denoted by $r$. Landscape properties for which the correlation has a raw $p$-value < 0.01 are appended by *.

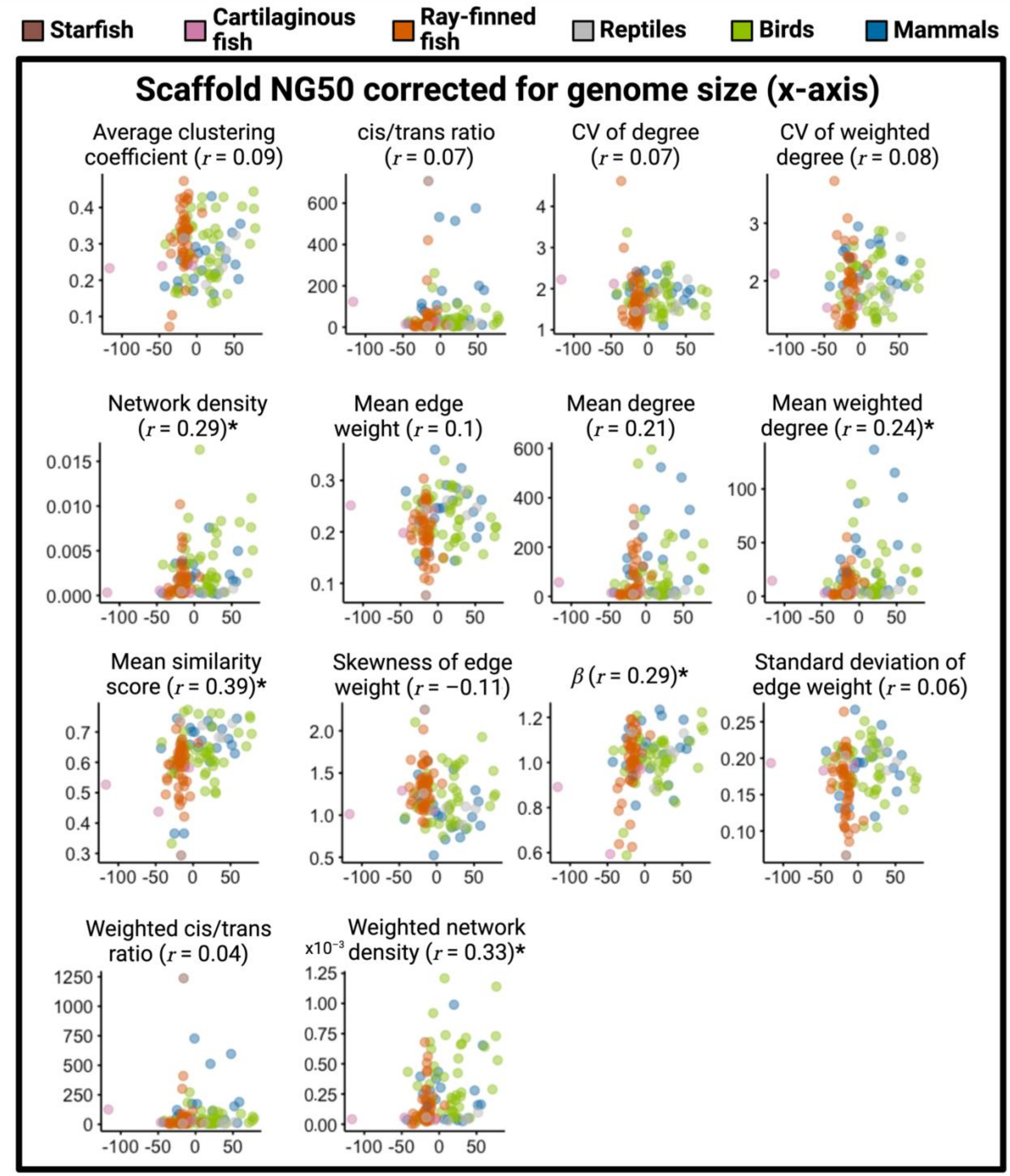


**Figure S6. The relationship between scaffold NG50 corrected for genome size and segmental duplication landscape metrics.** The horizontal axis represents the residuals from the linear model. The vertical axis represents the segmental duplication landscape property whose name is shown on the top of each figure panel. A dot represents a species, color-coded by taxonomy class shown in the legend. The Pearson correlation coefficient is denoted by $r$. Landscape properties for which the correlation has a raw $p$-value < 0.01 are appended by *.

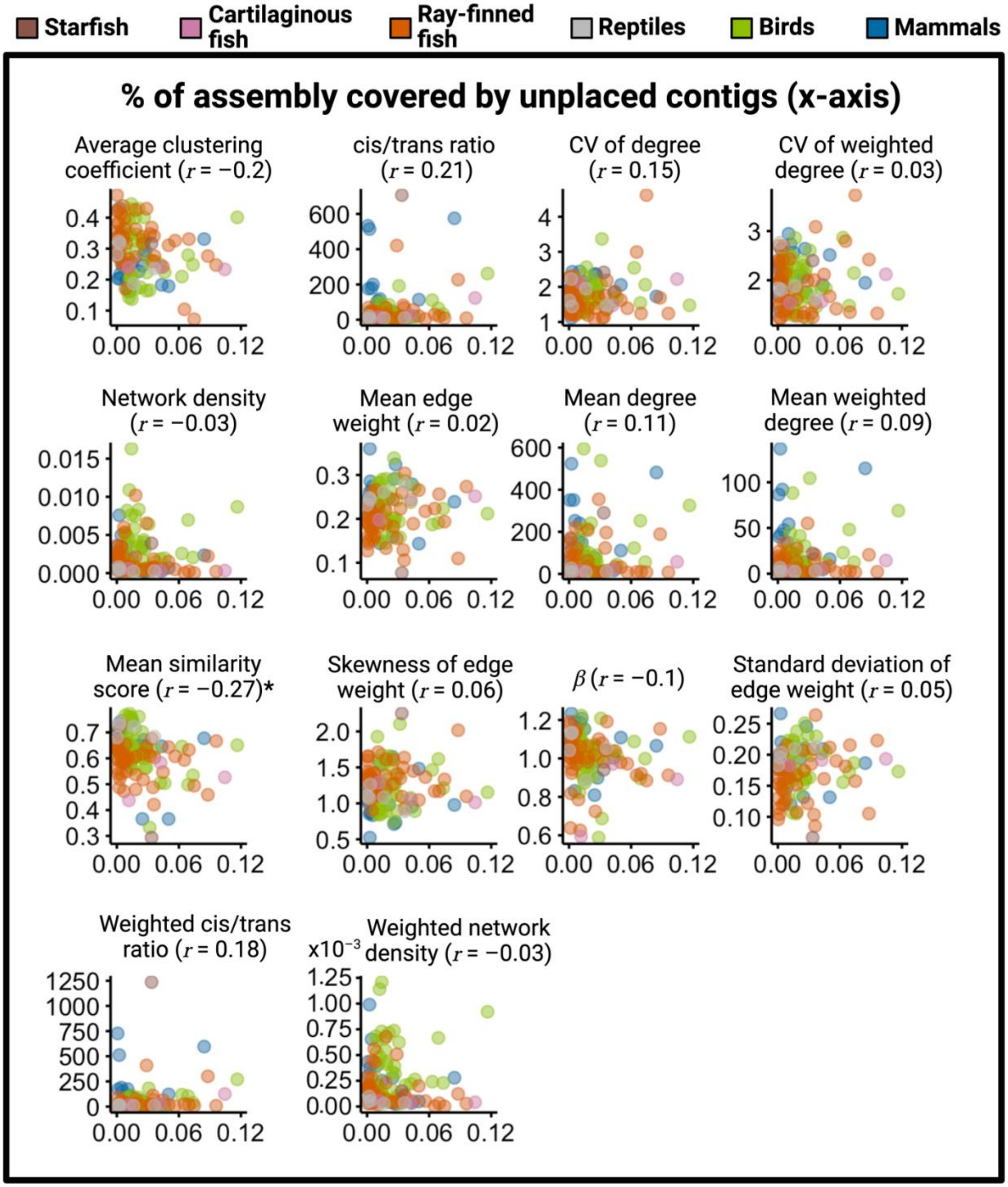


**Figure S7. The relationship between the percentage of genome assembly covered by unplaced contigs and segmental duplication landscape metrics.** The horizontal axis represents the unplaced contig coverage. The vertical axis represents the segmental duplication landscape property whose name is shown on the top of each figure panel. A dot represents a species, color-coded by taxonomy class shown in the legend. The Pearson correlation coefficient is denoted by $r$. Landscape properties for which the correlation has a raw $p$-value < 0.01 are appended by *.

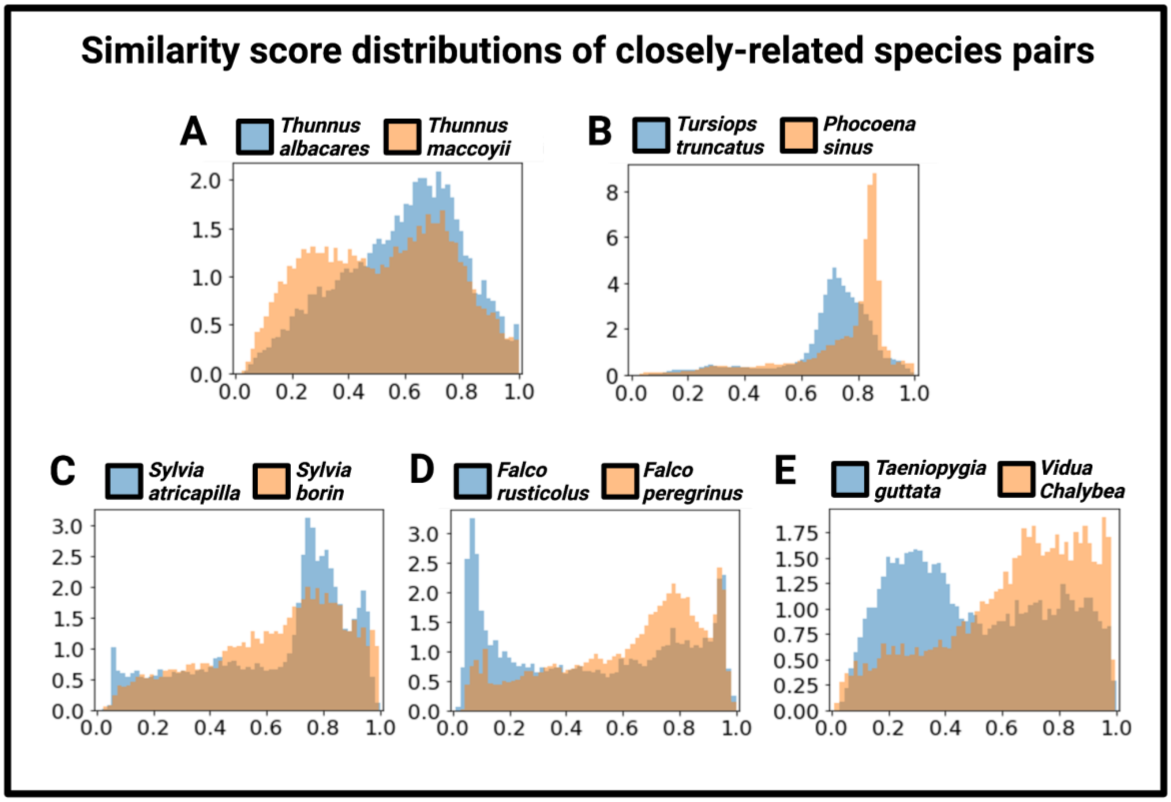


**Figure S8. Similarity score distributions for closely-related species pairs.** Five pairs of species that are phylogenetically close but have highly variable similarity distributions. The x-axis represents the similarity score and the y-axis represents density. **A.** Ray-finned fish. **B.** Mammals. **C-E.** Birds.

**Table S1. CV for each of the 14 measures across species.** For example, the "CV of node degree'', shown in the third row of the table, is a scalar measure calculated for each species. Its CV calculated based on all the ray-finned fish species is equal to 0.3427.

| | Ray-finned fish | Mammals | Birds | All |
|---|---|---|---|---|
| Mean degree | 1.3511 | 0.9542 | 1.4328 | 1.3952 |
| Mean node strength | 1.1969 | 1.0221 | 1.2969 | 1.4603 |
| CV of node degree | 0.3427 | 0.1585 | 0.2316 | 0.2689 |
| CV of node strength | 0.2945 | 0.1787 | 0.2259 | 0.2499 |
| Network density | 1.2066 | 0.9287 | 0.9382 | 1.1542 |
| Weighted density | 1.0628 | 0.9983 | 0.8133 | 1.0733 |
| Clustering coefficient | 0.2456 | 0.2653 | 0.3109 | 0.2841 |
| Density ratio (cis to trans) | 1.9270 | 1.1943 | 1.3387 | 1.9222 |
| Weighted density ratio (cis to trans) | 1.8682 | 1.2938 | 1.2629 | 2.2291 |
| $\beta$ | 0.1412 | 0.1075 | 0.1186 | 0.1316 |
| Mean edge weight | 0.2236 | 0.2236 | 0.1994 | 0.2316 |
| Std of edge weight | 0.2347 | 0.1927 | 0.1798 | 0.2159 |
| Skewness of edge weight | 0.1655 | 0.2667 | 0.2510 | 0.2385 |
| Mean similarity score | 0.1140 | 0.1598 | 0.1363 | 0.1496 |

**Table S2. The list of species for which we use a close substitution as proxies in place of the original species.**

| **Original species name** | **Substitute species name** |
|---|---|
| *Asterias rubens* | *Leptasterias muelleri* |
| *Pristis pectinata* | *Pristis clavata* |
| *Hemiscyllium ocellatum* | *Chiloscyllium punctatum* |
| *Salmo trutta* | *Salmo salar* |
| *Cottoperca gobio* | *Cottoperca* |
| *Anableps anableps* | *Anableps* |
| *Canis lupus orion* | *Canis lupus* |
| *Hemiprocne comata* | *Hemiprocne* |
| *Ciconia maguari* | *Ciconia ciconia* |
| *Phoenicopterus ruber* | *Phoenicopterus* |
| *Merops nubicus* | *Merops apiaster* |
| *Pogoniulus pusillus* | *Pogoniulus* |
| *Aquila chrysaetos chrysaetos* | *Aquila chrysaetos* |
| *Gopherus evgoodei* | *Gopherus agassizii* |
| *Notolabrus celidotus* | *Not substitution found* |

**Table S3**. **The correlation coefficient between the phylogenetic distance between two species and the distance in terms of each of the 14 measures.** The corresponding $p$-value is shown in the parentheses.

| | Ray-finned fish | Mammals | Birds | All |
|---|---|---|---|---|
| Mean degree | 0 (0.9883) | −0.12 (0.900) | 0.02 (0.4989) | 0.02 (0.0641) |
| Mean node strength | 0 (0.9070) | −0.11 (0.1204) | −0.01 (0.7426) | 0.01 (0.5584) |
| CV of node degree | 0.20 (1.0863× $10^{-10}$) | −0.02 (0.8186) | −0.09 (0.0103) | 0.07 (1.4593× $10^{-08}$) |
| CV of node strength | 0.06 (0.0368) | −0.10 (0.1842) | 0.12 (0.0002) | 0.02 (0.0490) |
| Network density | −0.11 (0.0006) | −0.03 (0.7043) | 0.07 (0.0475) | −0.04 (0.0006) |
| Weighted density | −0.12 (0.0001) | −0.08 (0.2447) | 0.01 (0.6822) | −0.03 (0.0085) |
| Clustering coefficient | 0.35 (6.1552× $10^{-32}$) | −0.10 (0.2046) | 0.08 (0.0128) | 0.04 (0.0029) |
| Density ratio (cis to trans) | −0.02 (0.5592) | −0.13 (0.0670) | −0.08 (0.0178) | 0.18 (6.3287× $10^{-54}$) |
| Weighted density ratio (cis to trans) | 0.01 (0.7203) | −0.14 (0.0629) | −0.08 (0.0149) | 0.23 (2.0282× $10^{-84}$) |
| $\beta$ | 0.06 (0.0708) | 0.18 (0.0109) | 0.09 (0.0067) | 0.04 (0.0005) |
| Mean edge weight | −0.05 (0.1308) | 0.41 (3.5282× $10^{-16}$) | −0.08 (0.0163) | 0.12 (1.1336× $10^{-23}$) |
| Std of edge weight | −0.06 (0.0746) | 0.34 (1.1546× $10^{-16}$) | 0.02 (0.6459) | 0.11 (3.7016× $10^{-21}$) |
| Skewness of edge weight | 0.04 (0.2123) | 0.48 (2.9343× $10^{-12}$) | −0.09 (0.0065) | 0.12 (5.0953× $10^{-25}$) |
| Mean similarity score | −0.06 (0.0412) | 0.85 (2.2588× $10^{-53}$) | 0.01 (0.8447) | 0.18 (1.6309× $10^{-47}$) |

**Table S4.** Effect sizes (Cohen's $d$) and p-values from the Mann-Whitney U test comparing species pairs with phylogenetic divergence less than 100 million years to pairs with greater divergence.

| | Ray-finned fish | Mammals | Birds | All |
|---|---|---|---|---|
| Mean degree | 0.0108 (0.6688) | 0.0269 (0.6532) | −0.0103 (0.0218) | 0.0789 ($5.1899\times 10^{-12}$) |
| Mean node strength | −0.0221 (0.7859) | 0.0395 (0.4435) | −0.0083 (0.0012) | 0.1015 ($9.8273\times 10^{-39}$) |
| CV of node degree | 0.0501 (0.0681) | 0.0351(0.7788) | −0.0009 (0.0088) | −0.1275 ($1.5750\times 10^{-39}$) |
| CV of node strength | 0.0514 (0.1737) | 0.0152 (0.8508) | 0.0669 ($5.4327\times 10^{-10}$) | −0.1084 ($1.1182\times 10^{-19}$) |
| Network density | 0.0858 (0.7273) | 0.0497 (0.4979) | −0.0497 ($3.1993\times 10^{-05}$) | 0.1306 ($3.6272\times 10^{-87}$) |
| Weighted density | 0.0997 (0.2668) | 0.0595 (0.4162) | −0.0548 ($1.1159\times 10^{-06}$) | 0.1713 ($5.7651$x$\times 10^{-125}$) |
| Clustering coefficient | −0.0199 (0.3565) | 0.0558 (0.4022) | −0.0024 (0.9342) | 0.0007 (0.0357) |
| Density ratio (cis to trans) | -0.0535 (0.9065) | 0.0465 (0.6276) | 0.0301(0.0646) | −0.0841(0.0738) |
| Weighted density ratio (cis to trans) | −0.0474 (0.9843) | 0.0433 (0.4041) | 0.0219(0.2666) | −0.0958 (0.0002) |
| $\beta$ | −0.0556 (0.0371) | 0.0171 (0.6979) | −0.0312 (0.0536) | −0.0358 (0.0005) |
| Mean edge weight | 0.0679 (0.1105) | −0.0011 (0.7281) | −0.0065 (0.9372) | −0.0456 (0.0002) |
| Std of edge weight | 0.0631 (0.2097) | 0.0047 (0.9468) | −0.0012 (0.1727) | −0.1383 ($1.6960\times 10^{-39}$) |
| Skewness of edge weight | 0.0365 (0.1249) | 0.0078 (0.9221) | 0.0136 (0.0099) | 0.0265 (0.0763) |
| Mean similarity score | −0.0129 (0.4847) | 0.0503 (0.3211) | −0.0412 ($7.3731\times 10^{-05}$) | −0.0384 ($9.3814\times 10^{-10}$) |

#### Supplementary Text S1

In (Abdullaev et al. 2021), the authors examined segmental duplication networks of nine species. They found that the component size distribution of the network was more similar within the six mammal species than between the mammal and non-mammal species. We computed the property that they used (i.e., the ordered sequence of the component size of the segmental duplication network) for each of our 118 species and computed the so-called Bray-Curtis dissimilarity (BCD) between each pair of species using the same method as theirs (Abdullaev et al. 2021). Then, as we did for our 14 quantities in the main analysis, we calculated the Pearson correlation coefficient between the phylogenetic distance between pairs of species and the BCD. The correlation was $r = –0.18$ ($p = 5.1905 \times 10^{-09}$), $r = 0.23$ ($p = 0.0015$), $r = 0.05$ ($p = 0.1635$), and $r = 0.14$ ($p = 4.5435 \times 10^{-33}$) for ray-finned fish, mammals, birds, and all species, respectively. Although the *p* values are mostly small due to large sample sizes, $r$ (as a measure of the effect size) is small. This result contrasts with those in (Abdullaev et al. 2021), which suggested that the component size distribution has an evolutionary signal, i.e., it is more conserved within mammals than between mammals and non-mammals. There are at least two possible reasons for the difference between their results and ours. First, we mainly compared within each of the three clades and used as many as 118 species. In contrast, their mammal versus non-mammal comparison is only based on six mammal species and three non-mammal species. Furthermore, one of the three non-mammal species used was *Caenorhabditis elegans*, which is evolutionarily farther than among the three clades we used. Second, the two studies used different computational methods to construct segmental duplication networks. Across-species comparison of segmental duplication networks aiming to improve the accuracy of segmental duplication calls and explicitly considering gene orthologs warrant future work.
